# Heparanase, a host gene that potently restricts retrovirus transcription

**DOI:** 10.1101/2024.12.06.627204

**Authors:** Brandon Waxman, Kyle Salka, Uddhav Timilsina, Supawadee Umthong, Deepak Shukla, Spyridon Stavrou

## Abstract

Heparanase (HPSE) is a heterodimeric β-D-glucoronidase that is critical in mammalian cells for the enzymatic cleavage of membrane associated heparan sulfate moieties. Apart from its enzymatic function, HPSE has important non-enzymatic functions, which include transcriptional regulation, chromatin modification and modulation of various signaling pathways. Interestingly, while *HPSE* is an interferon stimulated gene, past reports have shown that it has proviral properties for many different viruses, including Herpes Simplex Virus 1, as it assists virus release from infected cells. Yet any antiviral functions associated with HPSE have not been described. Here we show that HPSE utilizes a hitherto unknown mechanism to restrict retroviruses, by targeting the step of proviral transcription. Moreover, we demonstrate that HPSE blocks transcription initiation by targeting the SP1 transcription factor. Finally, we illustrate that the antiretroviral effect of HPSE is independent of its enzymatic activity. This report describes a novel antiviral mechanism utilized by HPSE to inhibit retrovirus infection.

**Importance:** Heparanase (HPSE) has emerged as an important factor that has proviral functions for a number of viruses including herpes simplex virus and hepatitis C virus, by assisting in virus egress. However, HPSE is an interferon stimulated gene and thus is a part of the host antiviral defense. Nothing is known about the antiviral functions of HPSE. Here, we examine in depth the role of HPSE during retrovirus infection using two retroviruses, human immunodeficiency virus type 1 (HIV-1) and murine leukemia virus (MLV). In this report, we show that mouse, but not human, HPSE blocks retrovirus infection by targeting provirus transcription. HPSE sequesters the SP-1 transcription factor away from the proviral promoter thereby inhibiting transcription inhibition. In conclusion, our findings identify a novel antiviral function of HPSE and its potential role as an inhibitor of zoonotic transmission of retroviruses.

## Introduction

The *Retroviridae* family is a diverse family of viruses, which includes members that infect a variety of species and cause the development of immunodeficiencies and cancer in infected organisms [1]. Although the most prominent members of this virus family are the human pathogens human immunodeficiency virus type 1 and 2 (HIV-1 and -2), the study of simple retroviruses such as murine leukemia virus (MLV) and mouse mammary tumor virus (MMTV) have contributed critical information for a better understanding of host-retrovirus interactions [2, 3].

Retroviruses upon infecting the cell, establish chronic infections by integrating their genome into that of the host. The retroviral genome is flanked by repeated sequences known as the long terminal repeats (LTRs). The 5’ LTR acts as the viral promoter and along with host transcription factors and RNA polymerase II are essential for the production of viral RNAs [4]. The U3 region within the 5’LTR is critical for the binding of transcription factors that regulate viral RNA synthesis in the infected cells [5–7].

Because retroviruses constantly attack the host genome, they exert selective pressure on the host. In turn, host cells have developed extensive antiviral defense mechanisms. Among these, is the type I interferon response, which activates a wide array of genes known as interferon stimulated genes (ISGs) that encode host factors that potently inhibit viral infections [8]. While the antiviral function of some of these factors has been elucidated [e.g. APOBEC3 proteins, Bst2/tetherin [9, 10]], for most, their antiviral mechanism is currently unknown.

An ISG, whose antiviral function is unknown is Heparanase (HPSE). HPSE is a β-D glucoronidase, which is known to cleave heparan sulfate (HS) moieties from the cell surface and extracellular matrix [11, 12]. HPSE is a highly conserved gene found in organisms ranging from zebrafish to humans [13–16]. HPSE is initially translated as a preproenzyme, whose signal peptide is removed in the ER producing the proenzyme (proHPSE), which is enzymatically inactive [17]. Activation of HPSE occurs in the late endosomes via the cleavage of proHPSE by Cathepsin L to an enzymatically active heterodimer consisting of a 50 KDa and 8 KDa subunit [17]. Apart from its enzymatic function cleaving HS, HPSE exhibits nonenzymatic functions that include enhancement of AKT signaling, entry into the nucleus for the regulation of gene expression including changes in inflammatory gene expression utilizing a currently unknown mechanism [11, 18–25]. In recent years, HPSE has emerged as an important factor for viral infections, having a beneficial role to a diverse number of viruses including herpes simplex virus 1 (HSV-1) [19, 26], human papillomavirus (HPV) [27, 28], hepatitis C virus (HCV) [29] and others. For the aforementioned viruses, HPSE enhances the egress of nascent virions from infected cells by cleaving HS from the surface of the cells, thereby resulting in HS shedding and facilitating the release of nascent viral particles. Interestingly, while high throughput screens have suggested that HPSE, being an ISG, has antiviral functions [8, 30], no study has hitherto determined any HPSE-associated antiviral phenotype.

Here, we report a novel function of HPSE vis-à-vis retrovirus infection. We show that mouse HPSE (mHPSE), unlike the human ortholog (hHPSE), potently restricts a number of retroviruses by blocking virus production. Moreover, we demonstrate that mHPSE causes the reduction of viral RNA levels by targeting retrovirus transcription. Subsequently, we identify that mHPSE targets the transcription factor SP1 and thereby inhibits provirus transcription. We further illustrate that the enzymatic activity of mHPSE is dispensable for its antiretroviral function and map the C’ terminal domain as critical for blocking retrovirus infection. Taken together, we reveal a novel antiviral mechanism, by which HPSE modulates SP1, a transcription factor that is essential for retrovirus gene expression, thereby decreasing retrovirus RNA levels.

## Results

### Transcription regulation of *Hpse*

To determine whether mouse *Hpse* is expressed in leukocytes, cell populations that are naturally infected by retroviruses *in vivo*, we isolated peripheral blood mononuclear cells (PBMCs) from wild type (C57B/6J) mice and cell sorted for B cells, T cells and dendritic cells (DCs) [2, 31, 32].

Transcript levels of *Hpse* were determined by RT-qPCR. We found that *Hpse* RNA levels are high in mouse DCs and B cells, while in T cells *Hpse* is expressed at low yet detectable levels (Figure 1A). To elucidate the effect of retrovirus infection on *Hpse* transcript levels, we infected MutuDC1940 (immortalized mouse DC line [33]) with MLV (multiplicity of infection [MOI] 5) and at different time points, we collected RNA and performed RT-PCR. We found that MLV infection did not affect *Hpse* expression (Figure 1B). Human *HPSE* is an ISG [34], to determine the effect of type I IFN on mouse *Hpse*, we treated bone-marrow-derived macrophages and bone marrow-derived dendritic cells (BMDMs and BMDCs) with murine interferon β (IFN- β; 500U/ml). We then collected RNA at different time points and performed RT-PCR to determine *Hpse* RNA levels over time. We observed that IFN-β treatment led to the upregulation of *Hpse* mRNA levels over time (Figure 1C), in agreement with previous reports [34]. Thus, we concluded that *Hpse* is expressed in cell types naturally infected by retroviruses, is an ISG, but is not induced by MLV infection.

**Figure 1.**
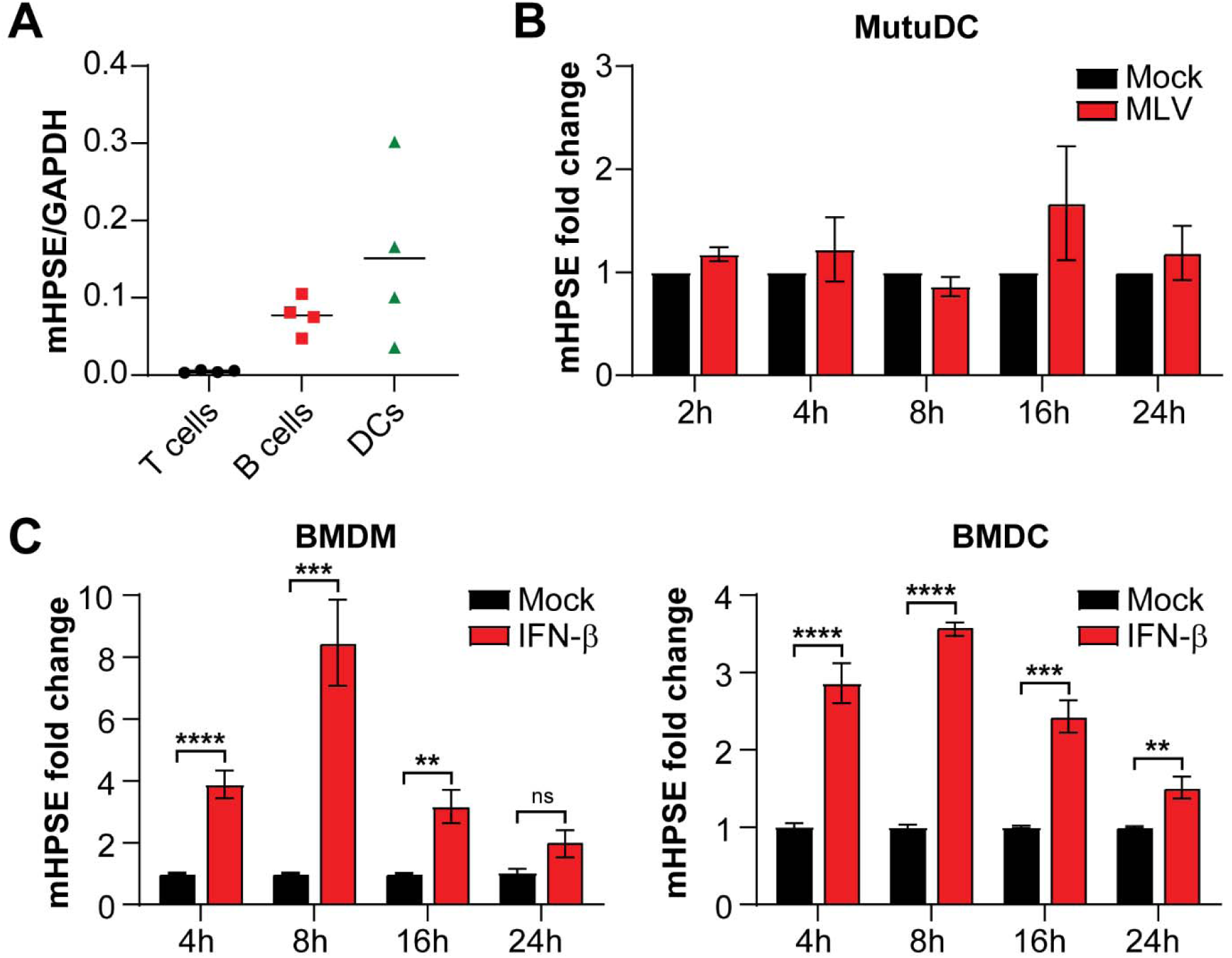
**Mouse *Hpse* is expressed in retrovirus target cells and induced by IFN-**β**. (A)** Peripheral blood mononuclear cells were collected from the blood of uninfected C57BL/6 mice and sorted for B cells, T cells, and dendritic cells, followed by RT-PCR to measure *Hpse* RNA levels. **(B)** MutuDC1940 cells were infected with MLV (5 MOI) and **(C)** Bone marrow-derived macrophages (BMDMs) and bone marrow-derived dendritic cells (BMDCs) were treated with 500U/ml of murine IFN-β. At the indicated time points post induction, *Hpse* RNA levels were determined by RT-PCR and fold change was calculated using the ΔΔCT method. Data presented as mean ± SEM. Statistical analysis was performed by multiple t tests, **(A)** N=4, **(B)** N=3, **(C)** N=6, ns; not significant, \*\**P*≤0.01, \*\*\**P*≤0.001, \*\*\*\**P*≤0.0001.

### Mouse Heparanase but not human Heparanase blocks retrovirus production

To determine the effect of the human and mouse HPSE orthologs on retroviruses, we co-transfected 293T cells with molecular clones for a number of retroviruses along with either hHPSE, mHPSE or an empty vector (EV). Cells and media were harvested 48 hours post transfection, followed by western blot assays to determine viral protein levels. To determine the effect of hHPSE and mHPSE on MLV, we utilized an infectious clone of Friend MLV [FMLV, pLRB302 [35]] and Moloney MLV [MMLV, p63.2 [36]], two ecotropic MLV strains. We found that for both strains of MLV, virus protein levels, both in the cellular and virus fractions, were reduced in the presence of mHPSE (Figure 2A). On the other hand, hHPSE had no effect on virus protein levels in either the cellular or virus fractions (Figure 2A). By using increasing amounts of mHPSE, we found that mHPSE reduced both the MLV envelope glycoprotein (gp70) and Gag (p65^gag^), in a dose-dependent manner (Figure 2B). Moreover, in agreement with previous studies [19, 37], we detected two forms of human and mouse HPSE in our transfected cells, a band at around 65 kDa representing the pro-enzyme form of HPSE (proHPSE, shown with an asterisk) and a band at 50 KDa representing the enzymatically active form of HPSE indicated with an arrowhead (Figure 2A). We also investigated the effect of human and mouse HPSE on another murine retrovirus, mouse mammary tumor virus (MMTV), by using a genetically engineered MMTV hybrid provirus (HP) clone [38]. Cells and virus-laden media were harvested followed by western blot assays. We found, similar to MLV, MMTV protein levels were reduced in the presence of mHPSE, while hHPSE had no effect (Figure 2C). Subsequently, we extended our studies to determine the effect of mouse and human HPSE on two lentiviruses, feline immunodeficiency virus (FIV) and human immunodeficiency virus type 1 (HIV-1). We co-transfected 293T cells with either an HIV-1 or an FIV infectious clone [NL4-3 and Petaluma [39] respectively] along with EV, mouse or human HPSE. Transfected cells were lysed 48 hours post transfection followed by western blots probing for HIV and FIV Gag. In the case of HIV-1, we also examined virus levels in the media of the transfected cells. Similar to what we observed with the murine retroviruses, only mHPSE reduced FIV and HIV-1 Gag levels, while hHPSE had no effect (Figure 2D and 2E). Because in the aforementioned experiments mHPSE caused the reduction of virus protein levels for all retroviruses tested, it is possible that mHPSE may reduce plasmid-mediated expression of proteins. To ensure that the effect of mHPSE on retrovirus protein levels is not due to interference with plasmid-mediated expression, we co-transfected 293T cells with either mHPSE or EV along with a GFP expressing plasmid and an MLV infectious clone. We observed that while mHPSE reduced MLV Gag protein levels, it had no effect on GFP protein levels (Figure 2F). Therefore, mHPSE does not interfere with plasmid-mediated translation and its effect is retrovirus-specific. In summary, our findings show that mHPSE reduces virus protein levels for a number of retroviruses, while hHPSE has no effect.

**Figure 2.**
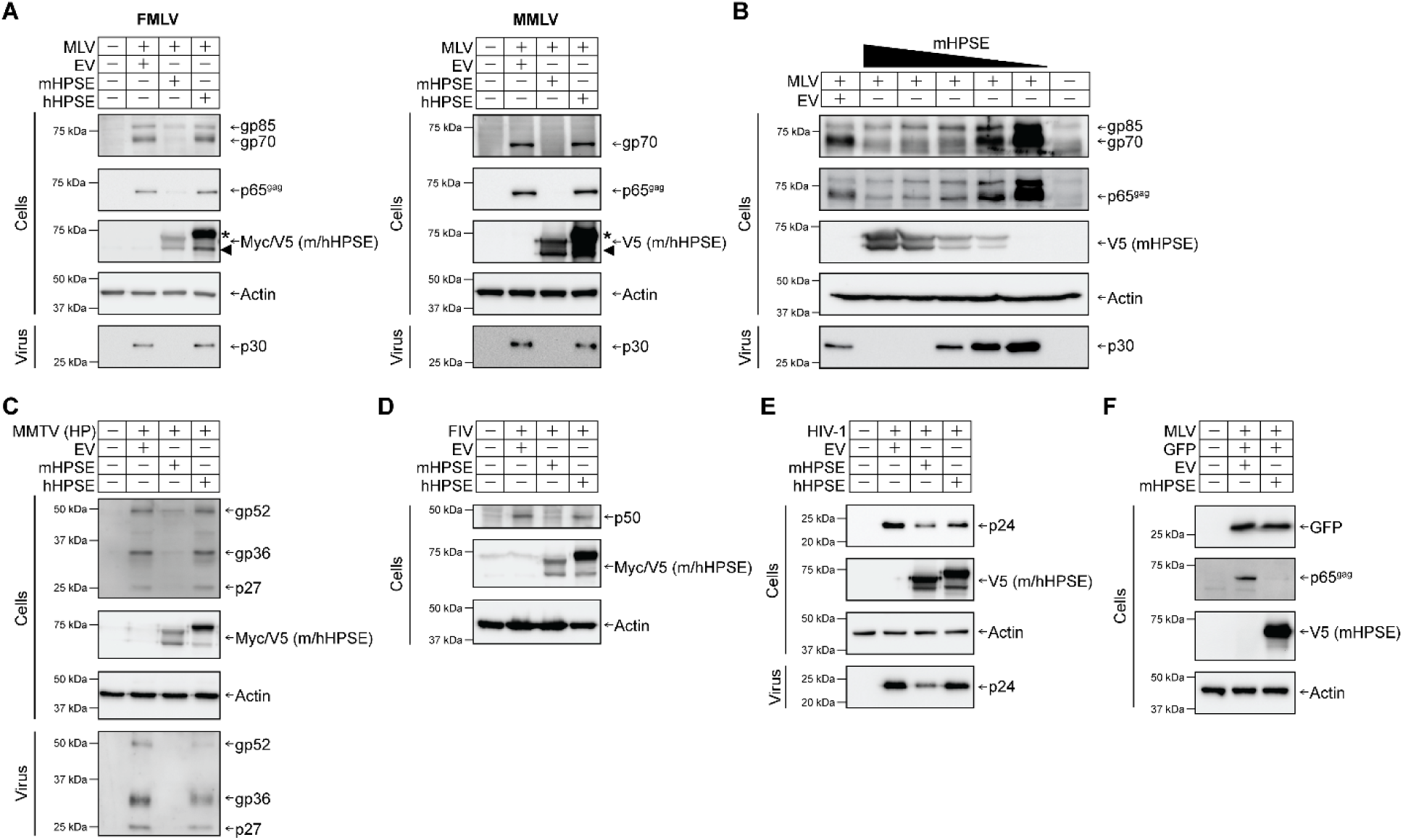
Mouse heparanase (mHPSE), but not human heparanase (hHPSE), is broadly antiretroviral. **(A)** mHPSE reduces protein levels of both Friend MLV (FMLV) (left panel) and Moloney MLV (MMLV) (right panel). **(B)** mHPSE reduces MLV levels in a dose dependent manner. mHPSE inhibits **(C)** MMTV, **(D)** FIV and **(E)** HIV-1. **(F)** mHPSE does not affect GFP protein levels. For **A** to **E**, 293T cells were co-transfected with plasmids encoding mHPSE, hHPSE, or empty vector (EV) along with infectious clones of FMLV, MMLV, an infectious genetically engineered MMTV hybrid provirus (HP), FIV or HIV-1. For **F**, 293T cells were co-transfected with plasmids encoding an FMLV infectious clone and a GFP expressing plasmid along with either mHPSE or EV constructs. Subsequently, cell lysate and virus-containing supernatant were harvested and samples were analyzed by immunoblotting for the indicated proteins. Representative immunoblotting results of 3 independent experiments are shown.

### Endogenous mHPSE restricts retrovirus infection

To determine the effect of endogenous mHPSE on retrovirus infection, we isolated mouse embryonic fibroblasts (MEFs) from 13-14.5 day embryos from C57BL/6J mice (wild type-WT mice) and Hpse knockout (Hpse KO) mice. Unfortunately there is no suitable antibody that detects endogenous mHPSE, thus we resorted to examining *Hpse* RNA levels in the HPSE WT and KO MEFs by RT-PCR (Figure 3A). Subsequently, cells were infected with MLV (0.02 MOI) and at 1, 2, 3 and 4 days post infection (dpi), DNA was isolated and qPCR was performed to determine virus DNA levels. We noticed that starting at 3 dpi MLV DNA levels in the *Hpse* KO MEFs were significantly higher (∼10x) compared to those seen in WT MEFs (Figure 3B). Thus, we concluded that endogenous mHPSE inhibits retrovirus infection.

**Figure 3.**
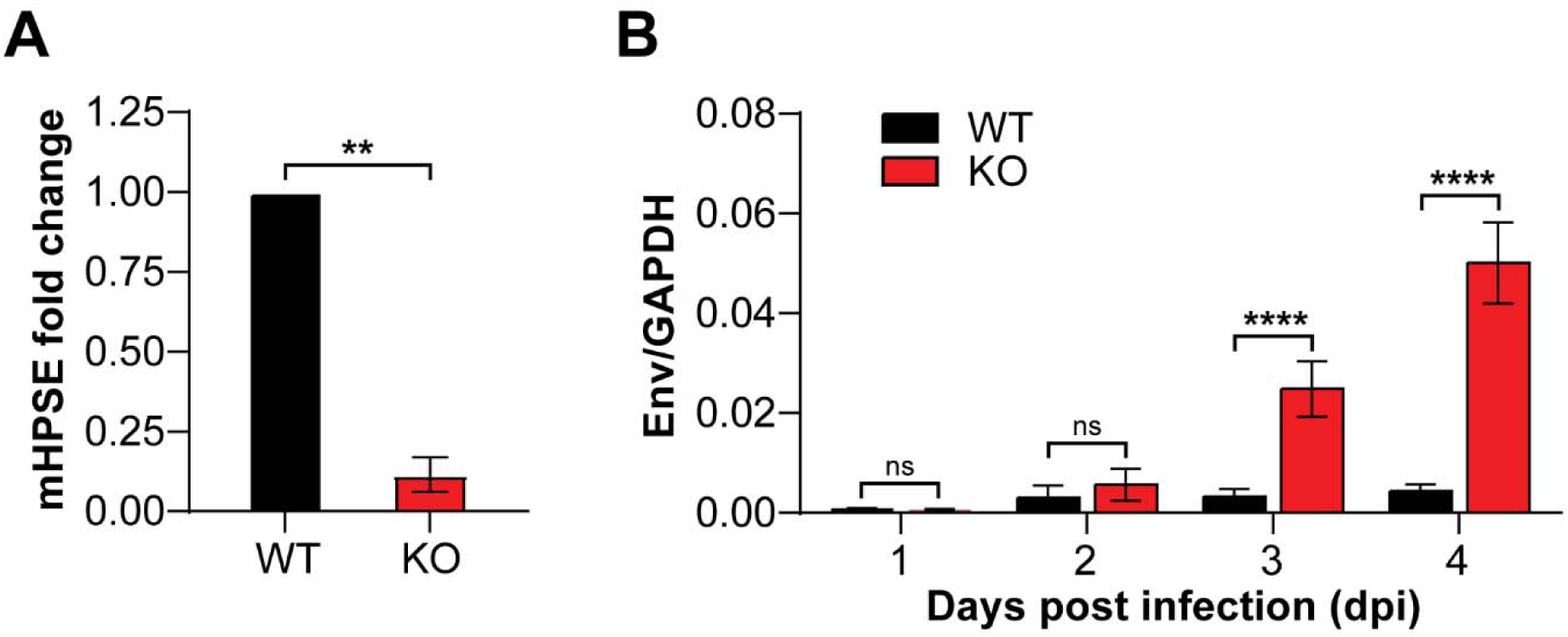
Endogenous mouse heparanase (mHPSE) restricts MLV infection. **(A)** Relative *mHPSE* RNA levels were determined by RT-PCR in mouse embryonic fibroblasts (MEFs) wild type (WT) and MEF mHPSE -/- (KO). **(B)** WT and KO cells were infected with MLV (0.02 MOI) and at the indicated time points post infection, MLV *env* DNA was measured by qPCR and normalized to mouse *Gapdh*. All data presented as mean ± SEM. Statistical analysis by **(A)** one sample t test or **(B)** multiple t tests, N=3, ns; not significant, \*\**P*≤0.01, \*\*\*\**P*≤0.0001.

### mHPSE inhibits retroviruses by targeting retrovirus transcription

Our aforementioned findings show that mHPSE is potently antiviral against a number of retroviruses (Figure 2). To better understand the mechanism by which mHPSE restricts retrovirus infection, we first investigated its subcellular localization. We transiently co-transfected 293T cells with either mHPSE alone or along with an MLV infectious clone. Whole cell, nuclear and cytoplasmic fractions were analyzed by western blots. We found that both pro- mHPSE (indicated with an asterisk-75 kDa) and enzymatically active form (indicated with an arrowhead-50 KDa) are present at high levels in the nuclear fractions, in agreement with previous reports [11, 18–20], and mHPSE nuclear localization is unaffected by the presence of MLV (Figure 4A).

**Figure 4.**
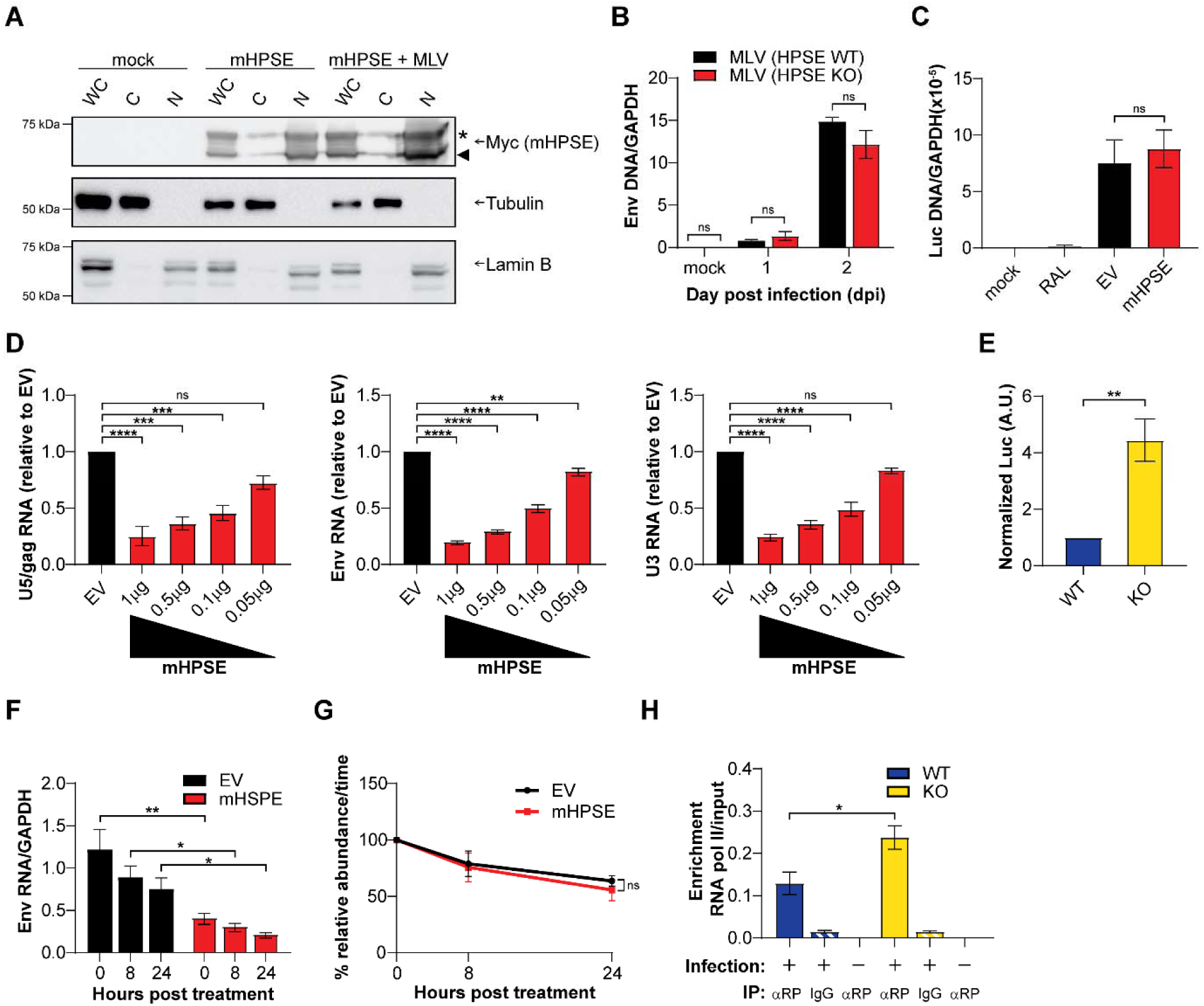
Mouse heparanase (mHPSE) restricts retrovirus transcription initiation. **(A)** mHPSE is primarily localized in the nucleus. 293T cells were transfected with plasmids encoding mHPSE or co-transfected with mHPSE and an MLV infectious clone, and then fractionated into whole cell lysate (WC), cytosolic fraction (C), or nuclear fraction (N). Samples were then analyzed by immunoblotting for the indicated proteins. Representative immunoblotting results of 3 independent experiments are shown. **(B)** mHPSE does not affect particle infectivity of progeny virions. Mouse embryonic fibroblasts (MEFs) wild type (WT) and MEF mHPSE -/- (KO) were infected with MLV (0.1 MOI), and equal amounts of the subsequently produced viruses were used to infect *Mus dunni* cells. At 24 hr post-infection, MLV DNA levels were determined by qPCR and normalized to *GADPH*. **(C)** mHPSE does not affect MLV integration. 293T-MCAT-1 cells were transfected with either mHPSE expressing or empty vector (EV) plasmids and then infected, in either the presence or absence of 100nM Raltegravir (RAL), with equal amounts of luciferase reporter pseudoviruses baring the MLV envelope. Viral DNA was measured by qPCR and normalized to *Gapdh*. **(D)** mHPSE reduces levels of all MLV viral transcripts. 293T cells were co-transfected with an MLV infectious clone and either EV or increasing amounts of mHPSE expressing plasmid. After 48 hr, MLV RNA transcripts were quantified by RT-qPCR with primers targeting the indicated regions. **(E)** Endogenous mHPSE restricts MLV expression. MEF WT and MEF KO cells were infected with equal amounts of MLV envelope pseudotyped luciferase reporter virus and at 2 dpi, luciferase expression was measured by luminescence assay and normalized to pseudovirus integration levels. **(F-G)** mHPSE does not affect the stability of MLV RNA. 293T cells were co-transfected with plasmids encoding MLV and either mHPSE or EV. Cells were then treated with 20 μg /ml of actinomycin D and MLV RNA levels were quantified by RT-qPCR at the indicated time points post treatment. **(F)** Data presented as mean MLV RNA normalized to *GAPDH*. **(G)** To measure the rate of RNA decay MLV RNA values from **F** were normalized to the 0 hr time. **(H)** mHPSE decreases the association of RNA pol II with the MLV LTR. MEF WT and MEF KO cells were infected with MLV envelope pseudotyped luciferase reporter virus. DNA was subsequently precipitated with an antibody targeting RNA pol II (αRP) or an isotype control (IgG), and enrichment of RNA pol II on the MLV LTR was determined by qPCR with primers targeting the MLV LTR and normalized to LTR DNA present in input samples. All data in graphs presented as mean ± SEM. Statistical analysis was performed by **(B,F)** two-way ANOVA, Sidak’s multiple comparisons test, **(C-D)** one-way ANOVA, Dunnett’s multiple comparisons test, or **(E, G, H)** unpaired t test, **(B-D)** N=3 and **(E-H)** N=4, ns; not significant, \**P*≤0.05, \*\**P*≤0.01, \*\*\**P*≤0.001, \*\*\*\**P*≤0.0001.

Initially, we investigated the effect of mHPSE on nascent viral particle intrinsic infectivity. Previous reports have shown that HS, the target of the enzymatic activity of mouse and human HPSE, can modulate entry for a number of viruses, including retroviruses, by affecting viral particle infectivity [40–43]. Therefore, it is possible that viruses produced in the presence or absence of mHPSE may display differences in viral particle intrinsic infectivity due to differences in HS moieties bound to the nascent virions. To determine the effect of mHPSE on viral particle infectivity, we infected Hpse WT and KO MEFs with MLV (0.1 MOI). Virus was harvested 2 dpi from the media of infected Hpse WT and KO MEFs and virus yields were determined by p30 (CA) ELISA. Subsequently, *Mus dunni* cells, which do not express endogenous mHPSE (Supplementary Figure S1), were infected with equal amounts of p30/CA of MLV collected from the media of either Hpse WT or KO MEFs and DNA was isolated from the infected *Mus dunni* cells 24 and 48 hours post infection followed by qPCR. We found that MLV DNA levels were similar in *Mus dunni* cells infected with virus derived either from Hpse WT or KO cells (Figure 4B). Thus, we concluded that mHPSE does not affect MLV intrinsic particle infectivity.

Subsequently, we decided to elucidate the step of the retroviral life cycle targeted by this host factor. Early events in the retrovirus life cycle include receptor binding, fusion, reverse transcription and integration of the reverse transcribed DNA into the host genome resulting in the creation of the provirus. Conversely, the late stages of the retrovirus lifecycle include transcription of the proviral DNA by the host RNA polymerase II, nuclear export of the viral RNAs, translation, assembly and release of the nascent virions. As mHPSE localizes in the nucleus (Figure 4A) and in our efforts to better understand the effect of mHPSE on the early events of retrovirus life cycle, we initially focused on the effect of mHPSE on retrovirus integration. To address this, we utilized 293T-MCAT-1 cells, 293T cells that are susceptible to MLV infection, as they constitutively express the ecotropic MLV receptor mouse cationic amino acid transporter 1 (mCAT-1) [44]. 293T-mCAT-1 cells were transfected with either a plasmid expressing mHPSE or EV. Afterwards, transfected cells were infected with equal amounts of MLV envelope pseudoviruses carrying a luciferase reporter and treated with or without Raltegravir (RAL;100nM), an integration inhibitor. As previously performed [45], cells were passaged for 6 days to eliminate any unintegrated viral DNA and thus any MLV DNA detected is from integrated proviral DNA. Subsequently, total DNA was isolated from the infected cells followed by qPCR. We found that integrated MLV DNA levels were unaffected by the presence or absence of mHPSE (Figure 4C). As expected, cells treated with RAL showed minimal levels of MLV DNA (Figure 4C). Therefore, we concluded that mHPSE does not affect retrovirus integration.

Another important step of the retrovirus life cycle that occurs in the nucleus is provirus transcription, which leads to the production of a number of retroviral transcripts that can either be translated to viral proteins or packaged as viral genomes in nascent virions. To examine the effect of mHPSE on viral RNA levels, we co-transfected 293T cells with increasing amounts of mHPSE along with an MLV infectious clone. RNA was isolated 48 hours post transfection and similar to a previous report [46], RT-qPCR was performed using three different sets of primers that detect initial (U5-gag), nearly (env) and fully (U3) completed MLV RNA transcripts. We found that all viral transcripts examined were reduced in the presence of mHPSE in a dose dependent manner (Figure 4D). As the aforementioned experiments were performed by overexpressing mHPSE, we also examined the effect of endogenous mHPSE on MLV expression levels. We infected Hpse WT and KO MEFs with equal amounts of MLV pseudoviruses carrying a luciferase reporter genome and 48 hours post infection we measured luciferase levels and normalized them to integrated MLV DNA levels. In agreement with our overexpression experiments, we found luciferase levels in Hpse KO MEFs were higher than those seen in WT (HPSE expressing) MEFs (Figure 4E). Therefore, we concluded that MLV expression levels are reduced in the presence of mHPSE. Yet, the decrease in retroviral RNA levels could be because of decreased transcription initiation, reduced RNA stability, nuclear export and splicing defects or a combination of the above processes. To determine the effect of mHPSE on RNA stability, 293T cells were co-transfected with an MLV infectious clone along with either mHPSE or EV. Cells were treated with actinomycin D (20 μg/mL) to inhibit transcription and total RNA was collected at different time points followed by RT-qPCR to measure for changes in MLV RNA levels over time in the presence or absence of mHPSE. As expected, we found that cells expressing mHPSE had significantly lower levels of MLV transcripts at all time points examined (Figure 4F). Nevertheless, the rate of RNA decay defined by the slope of relative MLV RNA abundance over time was similar between mHPSE expressing cells and those transfected with EV (Figure 4G). Thus, mHPSE does not cause the reduction of retroviral transcripts by modulating their stability. To assess the effect of mHPSE on retroviral transcript export from the nucleus, we co-transfected 293T cells with an infectious clone of MLV along with either mHPSE or EV. Transfected cells were harvested 48 hours post transfections followed by isolation of nuclear and cytoplasmic fractions using the PARIS™ kit (Ambion). Nuclear fraction purity was verified by western blot probing for the absence of GAPDH, a cytoplasmic marker (Supplementary Figure S2A). Subsequently, RNA was purified from the different fractions followed by RT-qPCR. We found that the ratio of nuclear over cytoplasmic viral RNA was similar in the presence or absence of mHPSE (Supplementary Figure S2B). Hence, we concluded that mHPSE does not affect the export of retroviral RNA from the nucleus. Afterwards, to determine the effect of mHPSE on the splicing of MLV transcripts, we co- transfected 293T cells with an MLV infectious clone and either mHPSE or EV. Cells were harvested 48 hours post transfection followed by RNA isolation and RT-qPCR using primers that allow for the discrimination between spliced and unspliced forms of MLV transcripts [47]. We found that mHPSE did not affect the ratio of unspliced to spliced MLV transcripts (Supplementary Figure S2C). Finally, we hypothesized that mHPSE inhibits provirus transcription initiation. To determine the effect of mHPSE on MLV transcription initiation, we infected WT and Hpse KO MEFs with MLV envelope pseudoviruses and performed a DNA pulldown assay precipitating for RNA Pol II followed by PCR with primers specific to the MLV LTR to measure the levels of RNA Pol II present on the viral LTR promoter in the presence or absence of endogenous mHPSE. We observed that there was a significant decrease in MLV LTR DNA levels precipitated with an antibody against RNA Pol II in the presence of mHPSE compared to EV (Figure 4H). Taken together, the above findings suggest that mHPSE exerts its antiretroviral effect by blocking provirus transcription initiation, thereby reducing viral RNA levels in infected cells.

### mHPSE inhibits retrovirus gene expression by targeting SP1

To further elucidate the mechanism by which mHPSE affects provirus transcription, we first investigated whether its effect is promoter-specific by determining the effect of mHPSE on luciferase expression levels under the regulation of various promoters. We first co-transfected 293T cells with a number of plasmids, in which the luciferase reporter gene is under the control of the indicated promoters [46], in the presence or absence of mHPSE. We found that mHPSE reduced luciferase levels regulated by the MLV and HIV-1 LTR as well as the HSV thymidine kinase (TK) promoter, but had no effect on the activity of the CMV immediate early (IE) and β- actin promoters (Figure 5A). As the above experiment was performed in 293T cells, a stable cell line of human origin, we then examined the effect of mHPSE in a murine cell line. We repeated the aforementioned experiment in NIH3T3 cells, a murine fibroblast cell line that does not express endogenous mHPSE (Supplementary Figure S1). Similar to our findings in 293T cells, we found that in NIH3T3 cells mHPSE reduced the activity of the MLV and HIV-1 LTR but not that of CMV IE promoter (Figure 5B). Thus, the effect of mHPSE on retroviral promoters is not cell line specific. To determine if endogenous mHPSE reduces the activity of the MLV and HIV- 1 LTR, we transfected Hpse WT and KO MEFs with the aforementioned luciferase promoter constructs. Luciferase levels were measured 24 hours post transfection and normalized to those under the control of the β-actin promoter. We found that luciferase levels under the control of the MLV and HIV-1 LTR were significantly higher in the Hpse KO MEFs when compared to the MEFs expressing mHPSE (WT MEFs) (Figure 5C). On the other hand, luciferase levels under the CMV IE promoter were unaffected by the presence or absence of endogenous mHPSE (Figure 5C). Thus, we concluded, that endogenous mHPSE specifically reduces the activity of the MLV and HIV-1 promoter. To determine the elements of the HIV-1 and MLV LTRs targeted by mHPSE, we initially utilized previously described HIV-1 LTR luciferase constructs containing mutations in their SP1, NFκB/NFAT, USF, TCF1α and NF-IL6 transcription factor binding sites [46, 48]. We transfected 293T cells with the different HIV-1 LTR constructs that contain mutations in the aforementioned transcription factor binding sites along with either mHPSE or EV followed by luciferase measurements 48 hours post transfection. Similar to previous reports, mutations in the SP1 binding site potently reduced the HIV-1 LTR activity (Supplementary Figure S3A) [46, 48]. In addition, we found that in the case of the HIV-1 LTR with mutations in the SP1 binding sites, mHPSE inhibited less potently both basal and Tat-mediated HIV-1 LTR gene expression when compared to the other HIV-1 LTR constructs with mutations in transcription factor binding sites (Figure 5D). Thus, we concluded that mHPSE inhibits HIV-1 LTR-mediated gene expression by targeting SP1. To determine if mHPSE also blocks MLV LTR-mediated gene expression by targeting SP1, we generated: (1) an MLV LTR luciferase reporter plasmid with mutations in six putative SP1 binding sites and (2) an MLV LTR luciferase reporter plasmid with mutations in the two CBF/Runx binding sites that served as our negative control (Supplementary Figure S3C). We co-transfected 293T cells with either the wild type or the two aforementioned mutant MLV LTR luciferase reporter plasmids in the presence or absence of mHPSE. We found that similar to our findings with the mutant HIV-1 LTR reporter constructs, the MLV LTR luciferase reporter plasmid with mutations in the SP1 binding sites had reduced LTR activity (Supplementary Figure S3B) and was less sensitive to mHPSE-mediated inhibition of MLV LTR gene expression (Figure 5E). To determine if SP1 is targeted by endogenous mHPSE, we transfected Hpse WT and KO MEFs with either the wild type MLV LTR luciferase reporter plasmid or the above described mutant MLV LTR luciferase reporter plasmids. Similar to what we saw in our overexpression experiment, endogenous mHPSE- mediated restriction was reduced in the MLV LTR reporter plasmid containing mutations in putative SP1 binding sites (Figure 5F). Over-expression of SP1 has previously been found to augment retrovirus production [46]. To determine if mHPSE suppresses such enhancement, we generated MLV carrying a luciferase reporter genome in the presence of increasing amounts of SP1 and either mHPSE or EV. Subsequently, we determined infectious virus yield by infecting NIH3T3 cells followed by measuring luciferase levels. We found that in the presence of mHPSE, over-expression of SP1 no longer enhanced MLV infectious virion production (Figure 5G).

**Figure 5.**
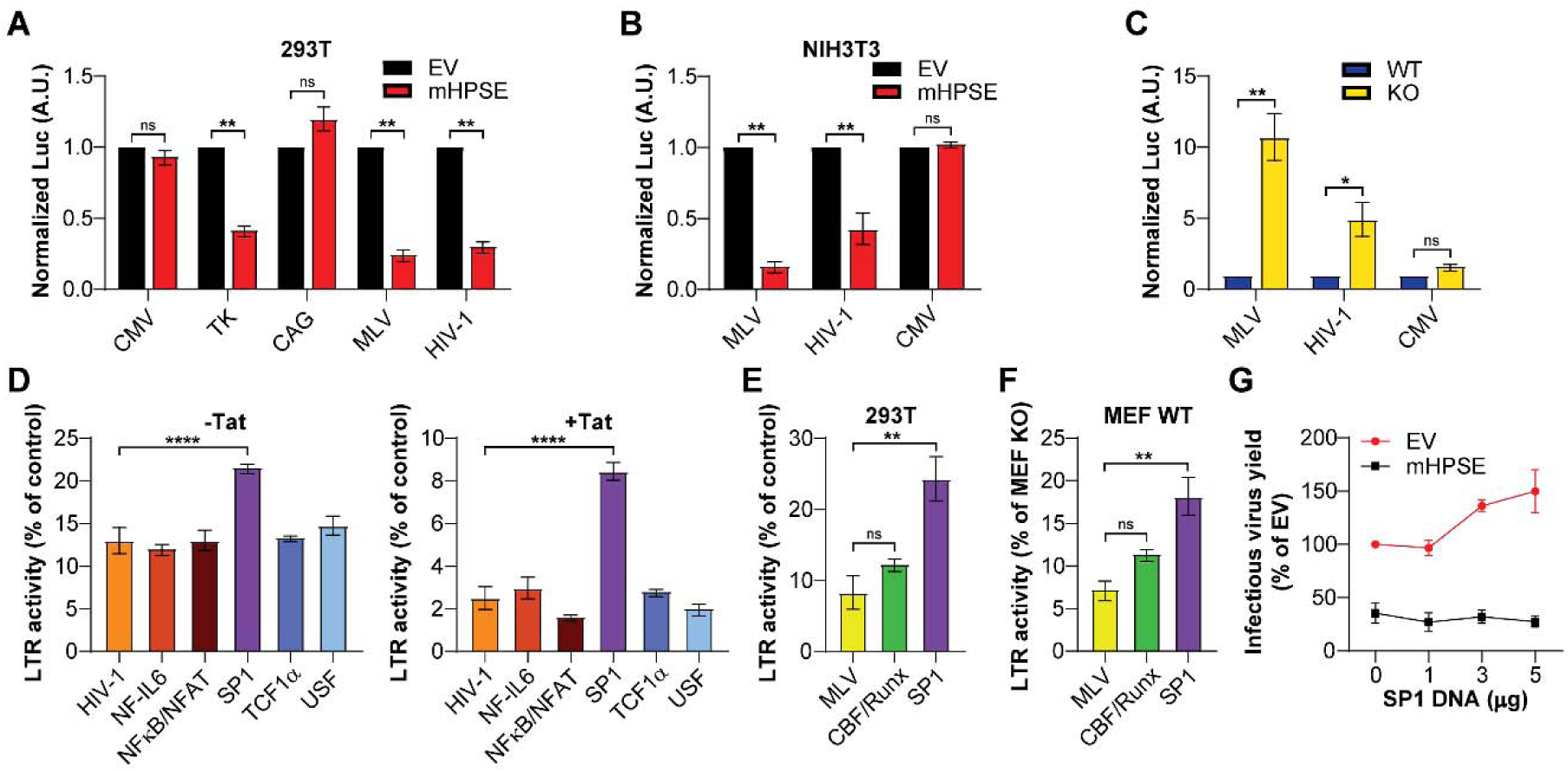
Mouse heparanase (mHPSE) targets SP1 to restrict retroviruses. (A-B) mHPSE restricts gene expression driven by the MLV LTR, HIV-1 LTR and the promoter of HSV thymidine kinase (TK) in **(A)** 293T and **(B)** NIH3T3 cells co-transfected with luciferase reporter constructs in which luciferase expression is driven by the indicated viral or cellular promoter, along with either empty vector (EV) or mHPSE expressing constructs. **(C)** Endogenous mHPSE restricts gene expression driven by both the MLV and HIV-1 LTR. Mouse embryonic fibroblasts (MEFs) wild type (WT) and MEF HPSE-/- (KO) cells were transfected with luciferase reporter constructs in which luciferase expression was driven by the MLV LTR, the HIV-1 LTR, and the CMV immediate early promoter (CMV) normalized to the luciferase activity of the β-actin promoter (CAG). **(D)** mHPSE targets SP1 to restrict HIV-1 expression. 293T cells were co- transfected with HIV-1 LTR luciferase reporter constructs containing the indicated transcription factor binding site mutations along with plasmids expressing either EV or mHPSE ± a plasmid encoding Tat. **(E-F)** mHPSE targets SP1 to restrict MLV expression. **(E)** 293T cells were co- transfected with MLV LTR luciferase reporter constructs containing mutations in the indicated transcription factor binding site in the presence of either EV or mHPSE. **(F)** MEF WT and KO cells were transfected with the same MLV LTR luciferase reporter constructs as in **E**. However, data presented in **F** shows MEF WT values normalized to those seen in MEF KO cells. **(G)** mHPSE inhibits SP1-mediated enhancement of MLV production. Luciferaser reporter MLV envelope pseudoviruses were produced in 293T cells in the presence of increasing amounts of SP1 along with either EV or mHPSE. Virus levels in the media of the transfected cells were then determined by infecting NIH3T3 cells and measuring luminescence. For panels **A-B** and **D-F**, data were normalized to samples lacking mHPSE. Data are all presented as mean ± SEM. Statistical analysis was performed by **(A-B)** one sample t test, **(C)** two-way ANOVA, Sidak’s multiple comparison test, or **(D, E, F)** one-way ANOVA, Dunnett’s multiple comparison test, **(A, B, E, G)** N=3 and **(C, D, F)** N=4, ns; not significant, \**P*≤0.05, \*\**P*≤0.01, \*\*\*\**P*≤0.0001.

To better understand the mechanism through which mHPSE targets SP1, we evaluated the effects of mHPSE on both endogenous murine and human SP1 protein levels, as both orthologs are about 90% identical in sequence. We transfected NIH3T3 and 293T cells with either EV or increasing concentrations of a plasmid encoding mHPSE and performed western blot analysis on cell lysate to determine the cellular levels of SP1. We found that both murine and human SP1 (mSP1 and hSP1) protein levels were unaffected by the presence of mHPSE in either NIH3T3 or 293T cells respectively (Figure 6A). Thus, we concluded that mHPSE does not lead to the degradation of either mouse or human SP1. However, it is possible that mHPSE binds mSP1 and sequesters it away from the 5’ LTR of the integrated provirus, thereby impacting provirus initiation. Initially, we investigated if mHPSE binds to mSP1. Unfortunately, the available antibodies for endogenous mHPSE are not suitable for co-immunoprecipitations (coIPs), thus we resorted to co-transfecting 293T cells with Myc-tagged mHPSE or mSP1 alone or mSP1 along with either myc-tagged mHPSE or hHPSE. CoIPs were then performed with either an antibody targeting mHPSE and hHPSE (αmyc) or an isotype control (mouse IgG), followed by western blot analysis. We found that only mHPSE co-immunoprecipitated mSP1 (Figure 6B right panel, lane 4). We then examined if the interaction between mSP1 and mHPSE is dependent on nucleic acids. We thus performed the aforementioned coIP in the presence or absence of benzonase, a nuclease that degrades both RNAs and DNAs. We observed that mHPSE precipitated higher levels of mSP1 in the absence of DNA, suggesting that DNA competes with mHPSE for binding to mSP1 (Figure 6C). To specifically examine the effect of mHPSE on the binding of mSP1 to the MLV LTR, we infected WT and Hpse KO MEFs with an MLV envelope pseudotyped reporter virus and performed DNA pull down using an SP1 specific antibody. MLV 5’ LTR levels were measured in the eluted DNA by qPCR using primers specific to the MLV 5’ LTR. We found that Hpse KO MEFs had higher levels of mSP1 bound to the MLV 5’ LTR as compared to WT MEFs, indicating that in the presence of endogenous mHPSE, there is a decrease in the mSP1 protein levels bound to the retroviral promoter (Figure 6D). Taken together, our findings indicate that mHPSE binds to mSP1, preventing it from associating with the 5’LTR of the MLV provirus, and thereby inhibiting transcription.

**Figure 6.**
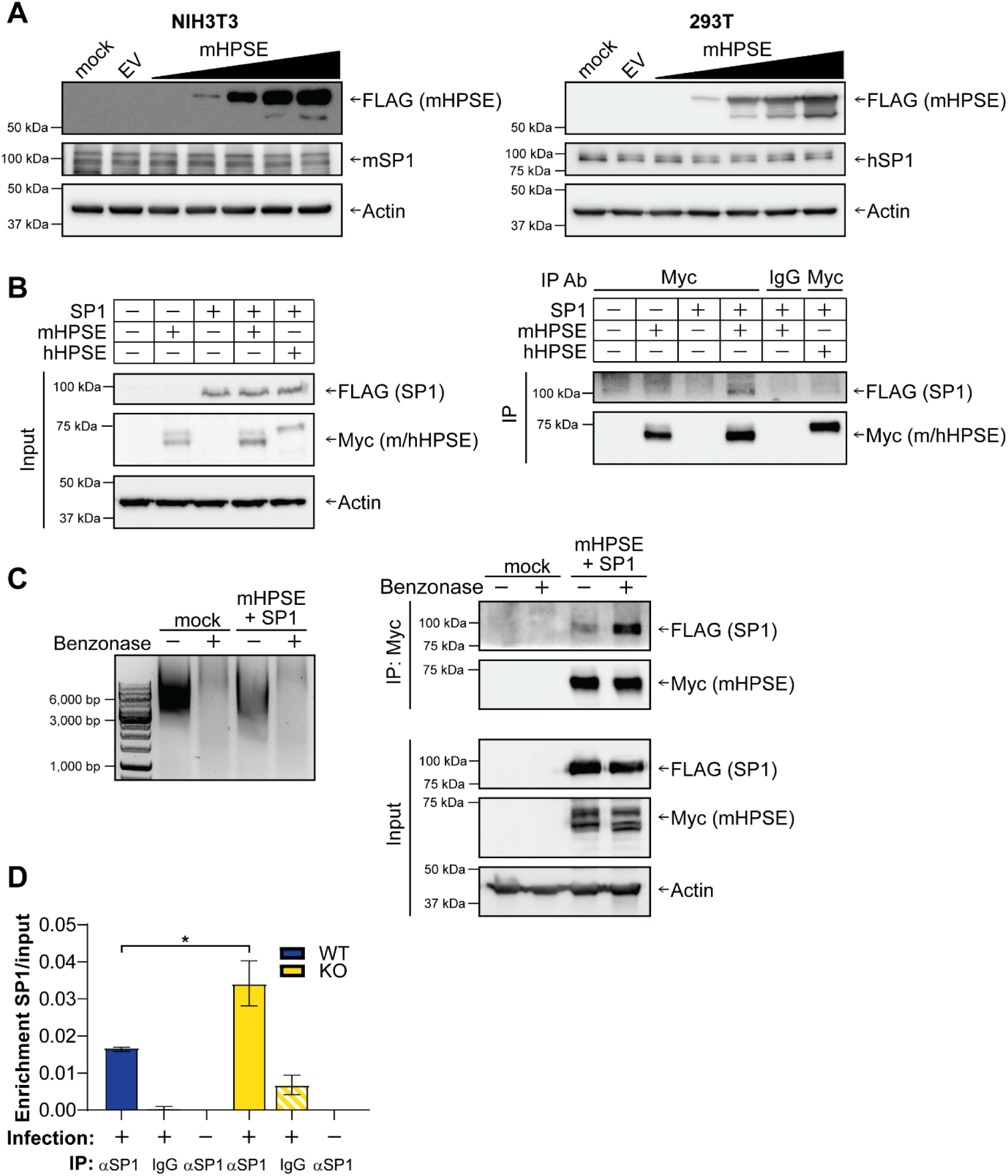
Mouse heparanase (mHPSE) sequesters SP1 to prevent it binding the retroviral LTR. **(A)** Neither mouse SP1 (mSP1) nor human SP1 (hSP1) are degraded by mHPSE. NIH3T3 (3T3, left panel) and 293T (right panel) cells were transfected with plasmids expressing either empty vector (EV) or increasing amounts of mHPSE and cell lysate was analyzed by immunoblotting. **(B-C)** mHPSE associates with SP1 and this interaction is independent of nucleic acids. 293T cells were either transfected solely with plasmids encoding EV, mHPSE, or SP1, or co-transfected with plasmids expressing SP1 and either mHPSE or human heparanase (hHPSE). Co-immunoprecipitation (coIP) was then performed either **(B)** without benzonase or **(C)** in the presence or absence of benzonase (750U/ml) using an anti-Myc (mHPSE) antibody or an isotype control (IgG). **(D)** mHPSE inhibits the association of SP1 with the MLV LTR. Mouse embryonice fibroblasts (MEFs) wild type (WT) and MEF HPSE -/- (KO) cells were infected with MLV envelope pseudotyped luciferase reporter viruses. DNA was subsequently precipitated with an antibody targeting SP1 (αSP1) or an isotype control (IgG), and enrichment of SP1 on the viral LTR was determined by qPCR with primers targeting the MLV LTR. MLV LTR DNA levels in pulldown samples normalized to input ± SEM are shown. Statistical analysis was performed by unpaired t test, N=3, \**P*≤0.05. For panels **A-C**, all samples were analyzed by immunoblotting for the indicated proteins and shown immunoblots are representative of 3 independent experiments.

### The nonenzymatic activity of mHPSE is critical for its antiretroviral function

Heparanase is known to have important enzymatic functions that are critical for its ability to break down HS chains, which in turn can be quite beneficial to virus infections [49]. In the case of HCV and HSV-1, the enzymatic activity of HPSE is crucial for the release of surface bound nascent viral particles [29, 50]. On the other hand, a number of HPSE nonenzymatic functions have been described revolving around cell signaling cascades (ERK, Akt etc), while regulation of gene expression by HPSE can be affected by both its enzymatic and nonenzymatic [51–53]. Yet, the role of the enzymatic and nonenzymatic functions of mHPSE in relation to its antiretroviral function is currently unknown. To determine the contribution of the enzymatic and nonenzymatic functions of mHPSE on its antiretroviral function, we first generated a mutant construct of mHPSE, in which the residues responsible for its catalytic activity [E217, proton donor and E335, nucleophile [54]] were mutated to alanines (mHPSE EE217/335AA). To verify that the mutations in the enzymatic active site do not disrupt the subcellular localization of mHPSE, we performed cell fractionations followed by western blot assays. We observed that mutating EE217/335 did not affect mHPSE subcellular localization (Supplementary Figure S4A). Subsequently, to ensure that the enzymatic activity of mHPSE EE217/335AA has been abrogated, we performed a HPSE enzymatic activity assay (Takara) of cell lysates from 293T cells transfected with either EV, WT mHPSE or mHPSE EE217/335AA and found that the enzymatic activity of mHPSE EE217/335AA was diminished (Figure 7A). Next, we measured luciferase levels using 293T cells transfected with plasmids encoding firefly luciferase under the control of either the HIV-1 or MLV LTR promoter (HIV-1 LTR-Luc and MLV LTR-Luc) in the presence of WT or enzymatically inactive (EE217/335AA) mHPSE or EV. We found that both WT and enzymatically inactive (EE217/335AA) mHPSE led to a similar reduction in luciferase reporter levels (Figure 7B). We also generated a mHPSE mutant, in which one of the HS binding sites (K150/153) was mutated to alanines (KK150/153AA). After verifying that the KK150/153AA mutation did not alter the subcellular localization of mHPSE (Supplementary Figure S4A), we performed a HPSE enzymatic activity assay and found the enzymatic activity of mHPSE KK150/153AA was intact (Supplementary Figure S4B). We attribute this to the fact that mHPSE has multiple HS binding sites [55], thus the loss of one of these site does not disrupt its enzymatic activity. Finally, when we performed luciferase reporter assays using the HIV-1 and MLV luciferase reporter plasmids mentioned above in the presence of either WT or KK150/153AA mHPSE, we found that luciferase levels were reduced under both WT and KK150/153AA mHPSE (Supplementary Figure S4C). Therefore the K150/153 HS binding site does not affect the antiretroviral functions of mHPSE.

**Figure 7.**
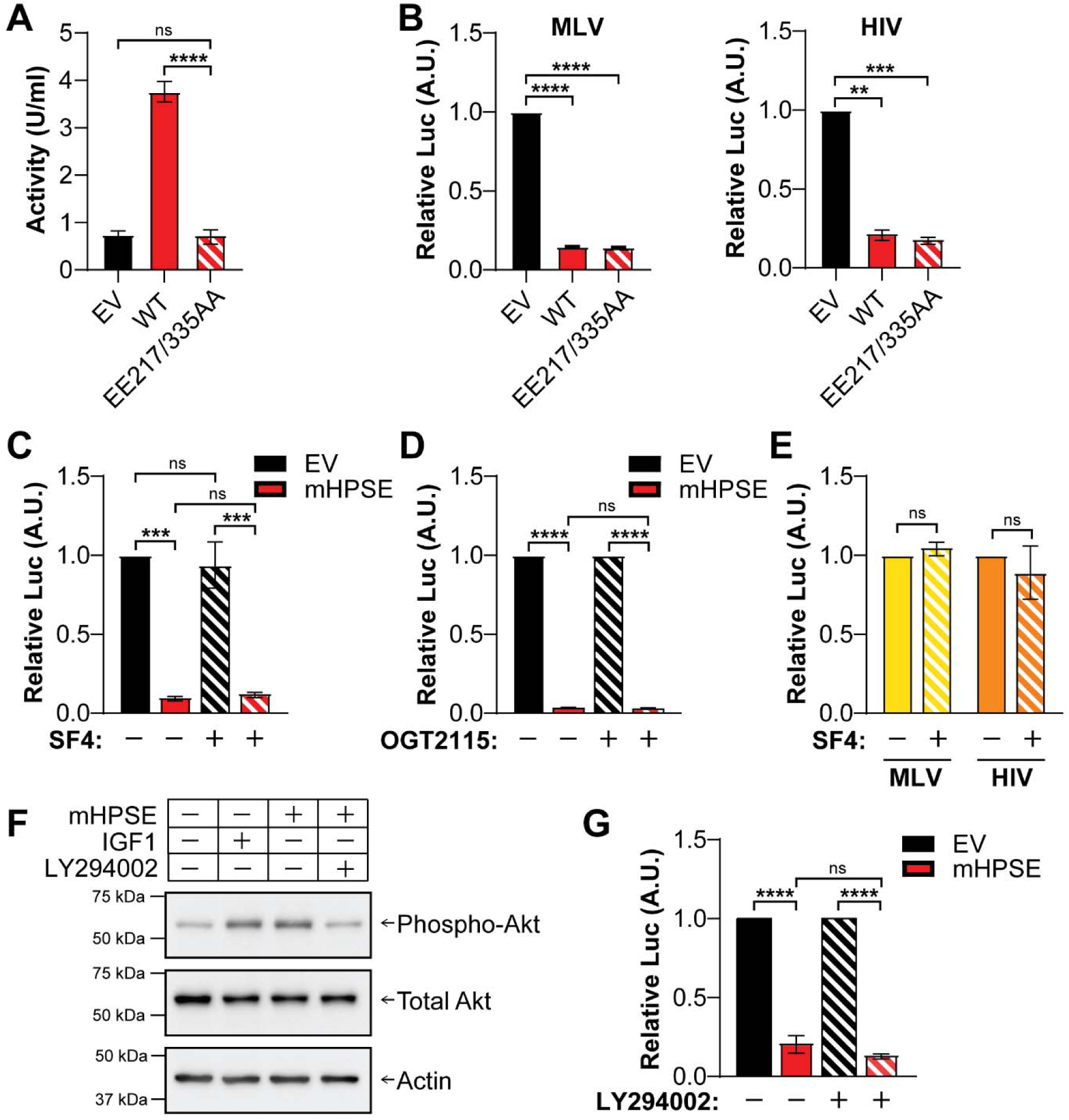
Mouse heparanase (mHPSE) restricts retrovirus expression through its nonenzymatic activity. **(A)** mHPSE glutamic acid residues 217 and 335 are required for enzymatic activity. 293T cells were transfected with plasmids encoding either empty vector (EV), mHPSE (WT), or mHPSE in which glutamic acid residues 217 and 335 are substituted for alanine (EE217/335AA) and enzymatic activity was measured by an ELISA-based absorbance assay. **(B-E)** The enzymatic activity of mHPSE is dispensable to its antiretroviral function. **(B)** 293T cells were co-transfected with MLV LTR or HIV-1 LTR luciferase reporter constructs and plasmids encoding either EV, mHPSE, or EE217/335AA. **(C-D)** 293T cells were co-transfected with plasmids encoding an MLV LTR luciferase reporter construct and either EV or mHPSE and then treated with **(C)** SF4 (100 μM), **(D)** OGT 2115 (10 μM), or vehicle control, followed by luminescence assays. **(E)** Endogenous mHPSE restricts retrovirus expression in the absence of enzymatic activity. MEF WT cells were transfected with either an MLV LTR or HIV-1 LTR luciferase reporter construct and treated with DMSO or SF4 (100 μM) followed by measurement of luciferase levels. **(F)** mHPSE increases activated AKT levels. 293T cells were treated with LY 294002 (10 μM) or vehicle control and then transfected with either EV, mHPSE, or insulin-like growth factor 1 (IGF1) constructs followed by immunoblotting. (G) mHPSE inhibits retrovirus expression through a mechanism distinct from its AKT signaling function. 293T cells were treated with LY 294002 (10 μM) followed by co-transfections with an MLV LTR luciferase reporter construct in the presence or absence of mHPSE followed by measurement of luciferase levels. For **F** representative immunoblotting results of 3 independent experiments are shown. For all luciferase reporter assays, luciferase expression was measured by luminescence, and values were normalized to EV or vehicle control samples. Data are all presented as mean ± SEM. Statistical analysis was performed by **(A, C, D, G)** one-way ANOVA, Tukey’s multiple comparison test, or **(B, E)** one sample t test, **(A)** N=4 or **(B-G)** N=3, ns; not significant, \*\**P*≤0.01, \*\*\**P*≤0.001, \*\*\*\**P*≤0.0001.

As the aforementioned studies were conducted using mutant constructs of mHPSE, we decided to complement our studies by using two commercially available pharmacological inhibitors of mHPSE enzymatic activity, heparastatin (SF4) and OGT 2115 [19, 56]. Both compounds block the enzymatic activity of mHPSE by binding to the catalytic pocket [57, 58]. 293T cells transfected with mHPSE were treated with either SF4 (100 μM) or DMSO and harvested 24 hours later. HPSE enzymatic activity assays were then performed to verify the inhibition of the enzymatic activity of mHPSE in the presence of SF4. In agreement with previous reports [56], we found that mHPSE enzymatic activity was diminished in the presence of SF4 (Supplementary Figure S4D). Subsequently, we transfected 293T cells with either mHPSE or EV along with a plasmid encoding firefly luciferase under the control of the MLV LTR promoter followed by treatment with either SF4 (100 μM) or DMSO. Luciferase levels were measured 24 hours later and similar to our results with the enzymatically inactive form of mHPSE, we found that the addition of SF4 did not affect the ability of mHPSE to decrease luciferase levels (Figure 7C). Similar to our findings with SF4, when we used another HPSE inhibitor, OGT 2115 (10 μM), we found that mHPSE reduced the activity of the MLV LTR both in the presence or absence of OGT 2115 (Figure 7D). In addition, we investigated the importance of the enzymatic activity of mHPSE, when it is expressed at endogenous levels. We used WT MEFs (mHPSE expressing MEFs) and treated them with DMSO or SF4 (100 μM) followed by transfection with a luciferase expressing plasmid under the control of the MLV or HIV-1 LTR promoters. We found that luciferase levels were similar in MEFs treated either with DMSO or SF4 (Figure 7E). Thus, we conclude that the enzymatic activity of mHPSE is dispensable for its antiretroviral effect.

Our data above demonstrate that mHPSE inhibits provirus transcription in a nonenzymatic manner. Yet human and mouse HPSE activate, via their nonenzymatic function, the AKT signaling pathway [11, 51, 59]. To ensure that mHPSE does not inhibit provirus transcription indirectly via the AKT signaling pathway, 293T cells were initially treated with the AKT inhibitor, LY 294002 (10 μM), followed by co-transfection with an MLV LTR luciferase reporter plasmid in the presence or absence of mHPSE and measurement of luciferase levels 24 hours later. Insulin growth factor 1 (IGF1) that is known to activate the AKT signaling pathway, served as positive control [60]. As expected, the presence of either IGF1 or mHPSE resulted in high levels of phosphorylated AKT while treatment with the AKT inhibitor reduced the levels of phosphorylated AKT to those seen in mock conditions (Figure 7F). When measuring luciferase levels, we found that mHPSE reduced MLV LTR luciferase activity the same regardless of the presence or absence of the AKT inhibitor (Figure 7G). Thus, we concluded that mHPSE does not inhibit provirus transcription via its activation of the AKT pathway, which agrees with our previous findings that show that mHPSE acts directly on provirus transcription. In summary, our findings show that the enzymatic activity of mHPSE and its effect on the AKT pathway are dispensable for the inhibition of retrovirus gene expression.

### The C’ terminal domain of the 50 KDa subunit of mHPSE is critical for its antiretroviral function

Human and mouse HPSE are heterodimers comprised of an 8 KDa and a 50 KDa subunit. The crystal structure of human HPSE has been solved and is composed of a (β/α)8-TIM barrel fold structure and a C’ terminal domain (aa 413-543) [61]. The TIM barrel fold is essential for its enzymatic activity and the C’ terminal domain, whose function is not well understood, is thought to be important for HPSE-mediated signaling and the non-enzymatic functions of HPSE [62]. To determine which domain of mHPSE is important for its antiretroviral function, we generated a number of chimeric HPSE molecules between mHPSE (has antiretroviral function-see Figure 2) and hHPSE (no antiretroviral function-see Figure 2). We initially constructed chimeras by swapping the 8 KDa subunits between human and mouse HPSE (Figure 8A-diagram); mHPSE^h8^, in which the 8 KDa subunit of mHPSE has been swapped with that of human HPSE and hHPSE^m8^, in which the 8 KDa subunit of human HPSE has been exchanged with that of mHPSE (Figure 8A). Unfortunately, hHPSE^m8^ was expressed at very low levels, possibly due to protein stability issues, thus we did not pursue this construct any further (Supplementary Figure S5). To confirm that swapping the 8 KDa subunit between the human and mouse HPSE, did not alter protein localization, we performed fractionation assays in 293T cells transfected with mHPSE^h8^. Subsequently, western blots were performed and we found that mHPSE^h8^ has similar subcellular localization, primarily in the nucleus and low amounts in the cytoplasm (Figure 8B), as that of WT mHPSE (Figure 4A). Thus, substituting the 8 KDa subunit of mHPSE with that of the human ortholog did not alter mHPSE localization. To determine the effect of mHPSE^h8^ on retrovirus restriction, we co-transfected 293T cells with an MLV infectious clone along with either EV, mHPSE, hHPSE or mHPSE^h8^. Media of the transfected cells were collected and cells were lysed 48 hours post transfectγion and western blots were performed to determine virus protein levels. We observed that chimeric mHPSE^h8^ resulted in a decrease of virus protein levels similar to what we found with WT mHPSE (Figure 8C). On the other hand, in agreement with our previous findings, virus protein levels were unaffected by hHPSE when compared to EV (Figure 8C). Thus, we concluded that the 8 KDa subunit of mHPSE is not important for its antiretroviral function and it is the 50 KDa subunit that is presumably responsible for the antiretroviral effect of mHPSE. Subsequently, we generated a new set of chimeric HPSEs, in which we replaced different regions of the mouse HPSE 50 KDa subunit with those from the 50 KDa subunit of the human ortholog; (a) m50K^hN^, the N’terminal region (aa 150-283) of the 50 KDa subunit of mHPSE is exchanged with those from hHPSE, (b) m50K^hM^, the 277-416 amino acids corresponding to the middle region of the 50 KDa subunit of mHPSE are substituted with those of hHPSE and c) m50K^hC^, in which the residues 410-535 found at the C’ terminal end of mHPSE are replaced with those from the same region in hHPSE and correspond approximately to the entire C’ terminal domain of mHPSE (Figure 8D). To verify that the chimeric HPSE molecules had similar subcellular localization as WT mHPSE, we transfected 293T cells with either m50K^hN^, m50K^hM^ or m50K^hC^, cells were then fractionated and processed for western blot analysis. We observed that the chimeric HPSE proteins showed similar subcellular localization as WT mHPSE (Figure 4A and 8E). To determine the effect of the different regions of the 50 KDa subunits on retrovirus restriction, we co-transfected 293T cells with an MLV molecular clone along with either wild type mHPSE or hHPSE or the m50K^hN^, m50K^hM^ or m50K^hC^ HPSE constructs we generated. Cells and media from the transfected cells were collected 48 hours post transfection. Cells were then lysed, while virus from the cell media was collected followed by western blot analysis. We found that while m50K^hN^ and m50K^hM^ restricted virus production similar to WT mHPSE, m50K^hC^ no longer inhibited virus production (Figure 8F). Therefore, we concluded that the C’ terminal region of the 50 KDa subunit of mHPSE is critical for its antiviral function. To determine if the C’ terminal region of mHPSE can confer an antiretroviral effect on hHPSE, we generated a chimeric hHPSE, in which its C’ terminal region is replaced with that of mHPSE, h50K^mC^ (Figure 8G). We co-transfected 293T cells with an infectious clone of MLV and either WT mHPSE, WT hHPSE or h50K^mC^. Cells and media were processed as before followed by western blot analysis. We observed that as before, WT mHPSE reduced virus protein levels, while hHPSE had no effect, interestingly, h50K^mC^ HPSE inhibited virus protein production similar to WT mHPSE (Figure 8H). In conclusion, we found that the C’ terminal domain of mHPSE is critical for its antiretroviral function.

**Figure 8.**
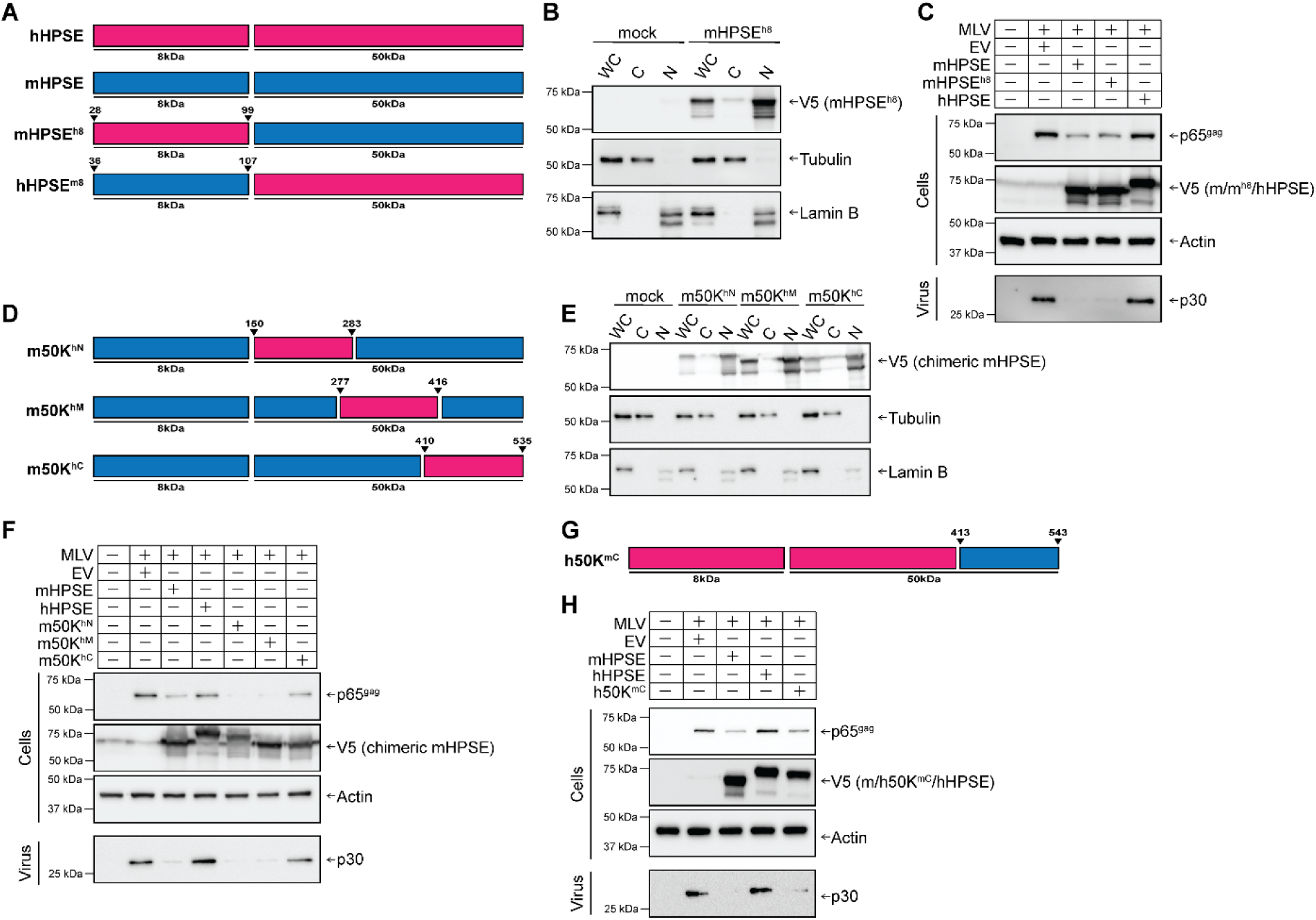
The C’ terminal domain of the mouse heparanase (mHPSE) 50 KDa subunit is responsible for MLV restriction. (A-C) The 8 KDa subunit of mHPSE is not responsible for its antiretroviral activity. **(A)** Diagram displaying the chimeric forms of mHPSE^h8^ and hHPSE^m8^ used in **B** and **C**. **(B)** The 8 KDa subunit of mHPSE does not affect its nuclear localization. 293T cells were transfected with plasmids expressing empty vector (EV) or mHPSE^h8^ and fractionated into whole cell lysate (WC), cytosolic fraction (C), or nuclear fraction (N) followed by immunobloting. **(C)** The 8 KDa subunit of mHPSE is dispensable for its antiviral effect. Assay was done the same as B, but an MLV infectious clone was included in the co-transfections. **(D- F)** The C’ terminal of the 50 KDa subunit of mHPSE is essential for retroviral restriction. **(D)** Diagram displaying the chimeric forms of mHPSE generated, m50K^hN^, m50K^hM^, and m50K^hC^. **(E)** m50K^hN^, m50K^hM^, and m50K^hC^ localize to the nucleus. 293T cells were transfected with either EV, m50K^hN^, m50K^hM^, or m50K^hC^ constructs and fractionated into WC, C, N followed by immunobloting. **(F)** Exchanging the C’ terminal domain of the 50 KDa subunit of mHPSE with that of hHPSE rescues MLV from mHPSE-mediated restriction. 293T cells were co-transfected with an infectious clone of MLV and either EV, mHPSE, hHPSE m50K^hN^, m50K^hM^, or m50K^hC^ followed by immunobloting. **(G-H)** The C’ terminal domain of the 50 KDa subunit of mHPSE confers the ability to restrict MLV onto hHPSE. **(G)** Diagram displaying the chimeric form of h50K^mC^ generated for this study. **(H)** Chimeric HPSE protein h50K^mC^ restricts MLV production. 293T cells were co-transfected with an infectious clone of MLV and either EV, mHPSE, hHPSE, or h50K^mC^ constructs followed by immunoblotting. Immunoblots shown in this figure are representative of 3 independent experiments.

## Discussion

Understanding host factors that target retrovirus transcription is of particular importance due to their implication in retrovirus latency and reactivation. A number of host factors that target retrovirus transcription have been identified over the years. Tripartite motif 22 (TRIM22) is an ISG that inhibits HIV-1 transcription in an E3 ubiquitin ligase independent manner and promotes latency [63, 64]. Another member of the TRIM family, TRIM28, suppresses HIV-1 expression and promotes latency by means of its SUMO E3 ligase activity and causes epigenetic modifications in the HIV-1 LTR promoter suppressing transcription [65]. In addition, interferon-γ inducible gene 16 (IFI16) and other pyrin and HIN domain (PYHIN) containing proteins also inhibit transcription in activated CD4+ T cells and macrophages [46, 66]. ZBTB2, a member of the POK (POZ and Kruppe) family of transcription factors, represses HIV-1 promoter by altering promoter histone acetylation [67]. Finally, the human silencing hub (HUSH) complex was recently shown to act as an important repressor of retrovirus transcription [68, 69].

In this report, we demonstrate that mHPSE has important antiretroviral functions. We found that mHPSE, similar to the human ortholog [70], is expressed at high levels in a number of different leukocyte populations including macrophages, B cells and DCs, cell types that are naturally infected by retroviruses [31, 71]. We also show mHPSE reduced virus production for a number of retroviruses including MLV, MMTV, FIV and HIV-1. The antiretroviral effect of mHPSE is particularly striking when compared to what is already known about the role of the human ortholog on viruses in general. All previous reports have demonstrated that hHPSE has in fact proviral function, as this host factor facilitates the release of nascent virions from infected cells for a number of viruses (e.g. HCV, HPV, HSV-1) by presumably cleaving HS found on the surface of the cells [29, 72]. The previous findings are quite paradoxical when taking into account that both HPSE orthologs are ISGs, and ISGs by their nature are meant to be a part of the host antiviral defense. On the other hand, large scale high throughput screens have suggested that HPSE has antiviral roles for a number of viruses [8, 30] and in this report we show that mHPSE is in fact antiretroviral. Surprisingly, our findings show that while mHPSE inhibits retrovirus production, hHPSE had no such effect. This difference in antiviral function between these two orthologs derived from different species is reminiscent of TRIM5α, a host gene that induces premature HIV-1 uncoating [73]. While human TRIM5α has a very weak antiviral effect against HIV-1, most Old World monkey TRIM5α proteins (e.g., rhesus macaque TRIM5α) restrict HIV-1 infection [74, 75]. Therefore, it is thought that TRIM5α has important protective function in cells against cross-species transmission of HIV-1 and other retroviruses [74]. In a similar manner, mouse cells are not permissive to HIV-1 infection due to numerous host factors that block virus replication. It is possible that mHPSE is one of those cellular factors that contribute to protecting murine cells from HIV-1 infection, something that needs to be further explored in the future. In addition, the difference in the ability of human and rhesus macaque TRIM5α to restrict HIV-1 has been attributed to the B30.2(SPRY) domain [76]. In a similar manner, we have identified that the C’ terminal domain of mHPSE is responsible for its antiviral effect. In fact, when comparing residues 413-543 of human HPSE, which correspond to its C’ terminal domain [61], and those that map to the C’ terminal domain of mouse HPSE (410-535), they only share 73% sequence homology. Future studies will focus on elucidating the specific residue(s) in the C’ terminal domain that are responsible for the difference in the antiviral phenotype and a better understanding of the evolutionary relationship between these two HPSE orthologs.

We also elucidated that mHPSE blocks retrovirus infection by targeting provirus transcription in both human and murine cells. We found that in the presence of mHPSE, retroviral RNA levels are reduced. In fact, using a variety of luciferase reporter constructs under the regulation of various viral promoters, we found that mHPSE inhibits MLV and HIV-1 LTR promoter activity, suggesting a role in the nucleus, which is in alignment with our fractionation assays, in which we detected mHPSE in the nuclear fraction. While HPSE has important functions in the extracellular matrix (ECM), previous reports have shown that human HPSE can also enter the nucleus by means of the chaperon heat shock protein 90 (Hsp90) [24]. In the nucleus, HPSE can modulate the expression of a number of genes such as those associated with glucose metabolism and inflammation [25]. HPSE is suggested to modulate gene transcription in a number of proposed mechanisms including the activation of histone acetyltransferases in the cell, binding directly to the DNA and modulating NFκB-mediated transcription, a well described transcription factor [77, 78]. Our data add to the functions of nuclear HPSE. We conclusively show that mHPSE in the nucleus has an additional role, that is inhibiting retrovirus transcription. Interestingly, our findings show that in the case of human and murine homologues of HPSE, both mature and pro- HPSE are present in the nucleus, which is in line with previous reports [21, 62]. Previous studies have shown that cleavage of HPSE by cathepsin L in the lysosome results in its conversion from proHPSE to mature HPSE [79]. However, other studies have found that proHPSE can enter the nucleus, where processing can also occur, in a hitherto uncharacterized mechanism [21, 62]. We attempted to elucidate which form of mHPSE is responsible for the antiretroviral function observed (proHPSE vs mature HPSE) as they are both found in the nucleus and contain the C’ terminal domain, important for the antiretroviral function of mHPSE. However, due to the paucity of information regarding the mechanism by which mHPSE is processed in the nucleus, we have been unsuccessful in elucidating which form exerts its antiviral function. Future experiments will focus in determining the form of mHPSE responsible for its antiretroviral function.

Our data show that MLV and HIV-1 LTR promoters with the SP1 binding sites mutated were less sensitive to mHPSE-mediated inhibition of LTR-driven gene expression in both human and murine cells suggesting that the antiviral effect of mHPSE is dependent on SP1. In addition, we show that mHPSE associates with SP1 and that mHPSE competes with DNA in binding to SP1. Finally, we demonstrate that in the presence of mHPSE, lower levels of SP1 are associated with the MLV LTR. The above findings suggest that mHPSE sequesters SP1 thereby reducing the levels of SP1 that can bind to the 5’ LTR promoter, resulting in the reduction of retrovirus gene expression. SP1 is a transcription factor that interacts with the 5’ LTR core promoter of HIV-1 via three GC rich binding sites and NFκB to activate the RNApol II-dependent transcriptional machinery at the HIV-1 promoter [80]. Furthermore, SP1 binding sites in the 5’ LTR promoter are present in many retroviruses including MLV, HIV-1, and human T-lymphotrophic virus type 1 (HTLV-1) [81]. Interestingly, a number of host antiviral factors that target HIV-1 gene expression, have been shown to target SP1. IFI16 interacts with SP1, sequestering it, thereby reducing the levels of this transcription factor that can activate the HIV-1 LTR promoter [46]. TRIM22, an important inhibitor of HIV-1 gene expression, blocks SP1 from binding to the 5’LTR of the HIV-1 promoter [82]. Our findings show that mHPSE targets SP1 in order to block retrovirus gene expression. Because SP1 is used by so many disparate retroviruses, it may be the reason that so many host factors target it. SP1, during retrovirus gene expression, is also considered to be essential in recruiting the RNApol-II transcriptional machinery to the 5’ LTR promoter of HIV-1 and MLV [80]; this is further supported by the fact that three very different host factors (IFI16, mHPSE and TRIM22) target this transcription factor. Moreover, SP1 has been previously shown to be a limiting factor in HIV-1 transcription and can play important role in reactivation [46]. Our findings further emphasize the importance of SP1 and provide for future studies mHPSE as a model to study SP1 mediated retrovirus gene expression *in vivo*.

Both human and mouse HPSE, via their enzymatic function, cleave HS found on the surface of the cells and the ECM thereby altering a number of cellular processes. In this report, using a mutant form of mHPSE, in which its enzymatic activity has been ablated, and two drugs (SF4 and OGT 2115) that block the enzymatic activity of HPSE we demonstrate that mHPSE inhibits retrovirus gene expression in a non-enzymatic manner. Moreover, we further demonstrate that HPSE has a direct effect on retrovirus transcription as it binds to SP1 and it does not act on retrovirus gene expression indirectly through the AKT pathway, the main pathway activated via its nonenzymatic function. In short, we show that the enzymatic activity of mHPSE is dispensable for its antiretroviral effect, which is in contrast to its proviral role, which is dependent on its enzymatic activity.

In summary, we demonstrate a new antiviral factor, mHPSE, which is IFN-inducible and has broad antiretroviral effects by inhibiting retrovirus transcription. However, mHPSE can be defined as a “negative regulator”, similar to IFI16 and TRIM22 [46, 82], as it targets a host protein, SP1, that the virus co-opts to its benefit, while SP1 is also critical for a number of cellular functions [83].

## Materials and Methods

### Ethics statement

Housing and breeding of mice occurs according to the policies of the Institutional Animal Care and Use committee of the University at Buffalo. Animal use in this study was sanctioned by this committee (MIC 46038Y).

### Cells

Mouse embryonic fibroblasts (MEFs) were prepared by harvesting cells of 13-14.5 day old embryos from C57BL/6J wild type (WT) or HPSE knockout (HPSE KO) mice as previously described [84]. Embryonic tissue fibroblasts were dissociated in 0.25% Trypsin (Gibco). Cells were then resuspended in Dulbecco’s Modified Eagle Media (DMEM; Gibco) supplemented with 10% (vol/vol) fetal bovine serum (FBS; Sigma-Aldrich), 6 mM L-glutamine (Gibco), 100 μg/ml penicillin and streptomycin (P/S; Gibco), and 0.1mM β-Mercaptoethanol (β-ME; Bio Rad). Immortalization of MEFs was achieved through serial passaging the cells. *Mus dunni* cells (ATCC), 293T cells (ATCC), and NIH3T3 cells (ATCC) were all cultured in complete DMEM media containing 10% (vol/vol) FBS, 6 mM L-glutamine (Gibco), and 100 μg/ml P/S (Gibco). 293T-mCAT-1 cells [85] were cultured in DMEM supplemented with 8% (vol/vol) donor calf serum (DCS; Gibco) 6 mM L-glutamine (Gibco), 100 μg/ml P/S (Gibco), and 100 μg/ml Geneticin (Gibco). MutuDC1940 cells [33] were cultured in Iscove’s modified Dulbecco’s medium (IMDM) supplemented with 8% FBS, 100 mg/ml P/S, 1 mM sodium pyruvate, 10 mM HEPES (Corning), and 0.05 mM β-Me (Bio Rad). To generate BMDMs (bone marrow-derived macrophages) and BMDCs (bone marrow-derived dendritic cells) bone marrow was extracted from 6- to 9-week C57BL/6J mice and cells were isolated and cultured as previously described [86].

### Plasmids

The pCMV-Red Firefly Luc (CMV IE; 16156) and pTK-Red Firefly Luc (TK; 16157) constructs were purchased from Thermo Scientific, and pCAG-Luciferase (β-actin-Luc; 55764) and pGL3- MMLV-LTR-Luc (MLV-LTR-Luc; 67831) were purchased from Addgene. pGL3-HIV-1 LTR-Luc (HIV-1 LTR-Luc) and pGL3-HIV-1 LTR-Luc plasmids with mutations in SP1, NFκB/NFAT, USF, TCF1α or NF-IL6 transcription factor binding sites have been previously described [46, 48] and were obtained from Frank Kirchhoff. MLV-LTR-Luc constructs with mutations in SP1 binding sites or deletion of the CBF/Runx binding sites were designed by us and constructed by Genscript. Genetically engineered MMTV hybrid provirus [HP; [38]], Friend MLV [FMLV; pLRB302 [35]], Moloney MLV [MMLV; p63.2 [36]], FMLV Gag/Pol [87], GS3-hHPSE-V5/His [19], and hHPSE-V5/His [19] have been previously described. A mHPSE- Myc/FLAG expressing plasmid was purchased from Origene (MR221855). Empty vector (EV; pcDNA3.1Myc/his) was purchased from Invitrogen. Feline immunodeficiency virus (FIV; pFIV- 34TF10, NIH-ARP 1236) and human immunodeficiency virus type 1 (HIV; NL4-3, NIH-ARP 114) were both obtained from NIH AIDS Reagent program, Division of AIDS, NIAID. EGFP- N1 was purchased from Clontech. pFB-Luc was purchased from Agilent.

### Cloning

PSV MLV Eco was generated using the NEBuilder HiFi DNA Assembly Cloning Kit, as per the manufacturer’s recommendations. The vector backbone was first PCR amplified from pSV-A-MLV-env (NIH-ARP 1065) with pSV-FMLV-Vec-F and pSV-FMLV-Vec-R primers (Table S1) and FMLV envelope was PCR amplified from FMLV (pLRB302) with pSV-FMLV-env-F and pSV-FMLV-env-R primers (Table S1). The PCR products were then assembled into PSV MLV Eco. Mouse HPSE-V5/His was generated by inserting mHPSE into pCDNA3.1-V5/His (Invitrogen), by first introducing XbaI and HindIII restriction digestion sites at the 5’and 3’ends of mHPSE gene via PCR using mHPSEcloF and mHPSEcloR primers described in Table S1.

Insert and vector were digested with XbaI and HindIII followed by ligation. All additional cloning of mHPSE was performed using this pCDNA3.1-mHPSE-V5/His (mHPSE-V5/His) expression clone as a template. mHPSE^h8^(mHPSE containing aa 36-107 of hHPSE), hHPSE^m8^(hHPSE containing aa 28-99 of mHPSE), m50K^hN^(mHPSE aa 150-283 replaced with aa 158-290 of hHPSE), m50K^hM^(mHPSE aa 277-416 replaced with aa 282-424 of hHPSE), m50K^hC^(mHPSE aa 410-535 replaced with aa 418-543 of hHPSE) were generated using the NEBuilder HiFi DNA Assembly Cloning Kit as per the manufacturer’s recommendations.

Briefly, to produce mHPSE^h8^, mHPSE-V5/His was PCR amplified with MuHep_hu8KDF and MuHep_hu8KDR primers (Table S1) and aa 36-107 of hHPSE-V5/His was PCR amplified with hu8KD_m50F and hu8KD_m50R primers (Table S1) followed by assembly of PCR products. The inverse chimera, hHPSE^m8^, was made by first PCR amplifying aa 28-99 of mHPSE-V5/His with Mu8KD_h50F and Mu8KD_h50R primers (Table S1) and hHPSE-V5/His with HuHep_M8KDF and HuHep_M8KDR primers (Table S1) followed by assembly of PCR products. To generate m50K^hN^, mHPSE-V5/His was PCR amplified with CXm50^N^F and CXm50^N^R primers (Table S1) and hHPSE-V5/His was PCR amplified with CXh50^N^F and CXh50^N^R primers (Table S1). For the m50K^hM^ expression clone, mHPSE-V5/His was PCR amplified with CXm50^M^F and CXm50^M^R primers (Table S1) and hHPSE-V5/His was PCR amplified with CXh50^M^F and CXh50^M^R primers (Table S1), while for m50K^hC^ expression vector, PCR amplification of mHPSE-V5/His was performed using CXm50^C^F and CXm50^C^R primers (Table S1) and for hHPSE-V5/His the CXh50^C^F and CXh50^C^R primers were used (Table S1). The inverse construct, h50K^mC^(413-543 of hHPSE substituted with aa 405-535 of mHPSE), in which the C terminus of hHPSE-V5/His was replaced with that of mHPSE-V5/His, was constructed by PCR amplification of mHPSE-V5/His with the primers mCdomainF and mCdomainR (Table S1), and hHPSE-V5/His with the primers hΔCF and hΔCR (Table S1). For all constructs mentioned above, once PCR amplification was completed, assembly of PCR products took place using the aforementioned kit. Myc tags were added to the N-terminus of mHPSE-V5/His (at aa 27) and hHPSE-V5/His (at aa 3) downstream of the signal peptide, by PCR amplification of mHPSE-V5/His and hHPSE-V5/His, respectively, with the primers: mHPSE_Myc_N_term_F and mHPSE_Myc_N_term_R or hHPSE_Myc_N_term_F and hHPSE_Myc_N_term_R (Table S1). The glutamates at resides 217 and 335 in mHPSE-V5/His were mutated to alanines in mHPSE-V5/His EE217/335AA through sequential PCR amplification using the E335F and E335R primers and E217F and E217R primers (Table S1) followed by ligation. Lastly, the lysine residues at positions 150 and 153 in mHPSE-V5/His were mutated to alanines by PCR amplification with the KK150/153AA_F and KK150/153AA_R primers (Table S1), followed by T4 ligation.

### Luminescence assays

Luminescence assays were performed using the Steady-Glo luciferase assay system (Promega, 2510) per the manufacturer’s recommendation and luciferase levels were measured using a Biostack4 (BioTek) luminometer.

### Transcriptional regulation of mouse *Hpse*

*Hpse* transcript levels in murine B cells, T cells, and DCs were determined as described before [87]. Briefly, blood was collected via retro-orbital bleeding from adult C57BL/6N mice (6 weeks old) and PBMCs were harvested using Ficoll-Plaque PLUS (GE Healthcare). T cells, B cells and DCs were sorted using a BD FACSAria fusion cell sorter after staining PBMCs with anti-CD3-APC (anti-CD3-allophycocyanin) (BD Biosciences), anti-CD45R/B220-FITC (anti-CD45R/B220-fluorescein isothiocyanate) (BD Biosciences) or anti-CD11c-APC (BD Biosciences) respectively. Total RNA was isolated using RNeasy minikit or RNeasy Plus Micro kit (Qiagen) per the manufacturer’s instruction and RT-qPCR was performed using a Power SYBR green PCR master mix kit (Applied Biosystems, A25742) in a CFX384 Touch real-time PCR detection system (Bio-Rad). The following primers were used: *mHPSE* (5’- CTGTCCAACACCTTTGCAGC-3’/5’CACTCGGAGTTTGCTCCTGT-3’ and *mGAPDH* (5′- CCCCTTCATTGACCTCAACTACA-3′/5′-CGCTCCTGGAGGATGGTGAT-3′). The relative levels of amplification were quantified for each sample from standard curves generated using known quantities of DNA standard templates. To determine the effect of MLV infection on *mHpse* transcript levels, MutuDC1940 cells were infected with MLV, and *mHpse* was quantified as previously described [86]. To determine the effect of mouse interferon β (IFN- β) on *mHpse* transcript levels, BMDMs and BMDCs were treated with 500U/ml of murine IFN- β and *mHpse* was quantified as previously described [86].

### MLV virus preparation

293T cells were seeded at 5 x 10^5^ per well in a 6-well plate. The following day, cells were transfected using Lipofectamine 3000, as per the manufacturer’s recommendations, with 3.5 μg of FMLV. At 48 hr post infection, virus was collected from the media of the infected cells followed by centrifugation at 700 x g for 10 min at 4°C, filtering through a 0.45 μm filter and treated with DNase (Roche) at a concentration of 1 μg/ml for 30 min at 37°C.

### HPSE retrovirus co-tranfections

To determine the effect of HPSE on MMTV, co-transfections of 293T cells were performed using calcium phosphate precipitation. Briefly, 293T cells were seeded in 10cm plates (4x10^6^ cells/plate) and the following day co-transfected with 13 μg of MMTV HP, 50 ng of rat glycocorticoid receptor (RSVGR) construct and 3ug of plasmids expressing either mHPSE- Myc/FLAG, hHPSE-V5/His, or EV. All other transfections were performed using a Lipofectamine 3000 transfection kit (Invitrogen, L3000075) as per the manufacturer’s recommendations. Briefly, 293T cells were seeded on a 6-well plate (5 x 10^5^ cells/well) and the following day were co-transfected with either 3.5 μg of FMLV, 3.5 μg of MMLV, 5.8 μg FIV, or 3.5 μg HIV along with 1 μg of either mHPSE-V5/His, hHPSE-V5/His or EV. To study the effect of mHPSE on plasmid-mediated transcription, 293T cells were seeded as described above and co-transfected with 2 μg of FMLV, 0.1 μg of EGFP-N1 and 0.3ug of either EV or mHPSE- V5/His. For all aforementioned transfections, cells and culture media were harvested 48 h after transfection. Cells were lysed in RIPA buffer (150mM NaCl, 1% NP-40, 0.5% sodium deoxycholate, 0.1% SDS, 25mM Tris, pH 7.4, with Halt phosphatase and protease inhibitors), and viruses from the culture media were purified and stored following ultracentrifugation through a 30% sucrose cushion as previously described [86]. Cell lysates and purified virions were then processed by western blot analysis.

### Western blot analysis

Cell lysates or virus-containing supernatants were mixed with 1x sample loading buffer, incubated for 10 min at 100°C and then resolved on 10% sodium dodecyl sulfate polyacrylamide gels. Protein samples was transferred onto PVDF membrane (BioRad) for 2 hr at 0.4 Amps followed by incubation in 5% milk in TBST overnight at 4°C before being probed with the following antibodies: goat anti-MLV gp70 (71), rat anti-83A25 (ATCC), goat anti-MMTV polyclonal (72), rat anti-MLV transmembrane protein/p15E (clone 42/114; Kerafast), rabbit anti-DYKDDDDK (Cell Signaling Technology, 2368S), rabbit anti-myc (Cell Signaling Technology, 2272S), mouse anti-V5 (Invitrogen, 46-0705), rat anti-MLV p30 (R187, ATCC), mouse anti-FIV p24 (PAK3-2C1, NIH-ARP, 4814), mouse anti-HIV p24 (NIH-ARP, 4121), rabbit anti-green fluorescent protein (Invitrogen, A11122), monoclonal anti-β-actin (Sigma-Aldrich), rabbit anti- SP1 (Proteintech, 21962-1-AP), mouse anti-SP1 (Santa Cruz, sc-17824), rabbit anti-Akt (Cell Signaling Technology, 9272S), and rabbit anti-P-Akt (S473; Cell Signaling Technology, 9271S). Secondary antibodies used in our studies are the following: horseradish peroxidase (HRP)- conjugated anti-rabbit IgG (Cell Signaling Technology, 7074S), HRP-conjugated anti-rat IgG (Cell Signaling Technology, 7077S), HRP-conjugated anti-mouse IgG (EMD Millipore), and HRP-conjugated anti-goat (Sigma-Aldrich). Imaging was performed using the enhanced chemiluminescence detection kits Clarity and Clarity Max ECL (Bio-Rad).

### Restriction by endogenous mHPSE

WT and HPSE-/- (KO) MEFs were seeded at 3 x 10^4^ cells in 12-well plates. The following day, cells were infected with FMLV (0.02 MOI) via spinoculation as previously described [88].

Briefly, virus was added to the cells followed by centrifugation at 250 x g for 2 hr at 22°C. Subsequently, inoculum was removed and replaced with fresh media. At 1, 2, 3, and 4 dpi, DNA was isolated using the DNeasy blood and tissue kit (Qiagen) and qPCR was performed in a CFX384 Touch Real-Time PCR Detection System using Power SYBR green PCR master mix kit and the following primers: *MLV env* (5’-TACAGGGAGCTTACCAGGCA-3’/5’-GTTCCTATGCAGAGTCCCCG-3’) and *mGAPDH* (5’-CCCCTTCATTGACCTCAACTACA- 3’/5’-CGCTCCTGGAGGATGGTGAT-3’).

### Cellular fractionation

Cells were collected in cold 1xPBS followed by centrifugation at 9,500 x g for 10 min at 4°C. The cellular pellet was then resuspended in PBS containing 0.05% Triton X-100 followed by the collection of an aliquot representing the whole cell fraction. The remaining resuspended cell lysates were incubated on ice for 30 min for lysis of the cell membranes followed by centrifugation at 2,400 x g for 5 min at 4°C to separate nuclear and cytosolic fractions, which were found in the pellet and supernatant, respectively. The cytosolic fraction-containing supernatant, was removed and saved and the nuclear fraction-containing pellet was washed 3 times in cold 1xPBS containing 0.05% Triton X100. All samples were then processed for western blot analysis (see “Western blots”). Fraction purity was confirmed by probing with mouse anti-Lamin B1 (Santa Cruz Biotechnology, sc-374015) and rabbit anti-β-tubulin (Cell Signaling Technology, 2128S).

### Mapping the antiretroviral domain of mHPSE

293T cells were seeded at 5 x 10^5^ cells per well in a 6-well plate and the following day transfected using Lipofectamine 3000, as per the manufacturer’s recommendations, with 1 μg of either EV, mHPSE or hHPSE along with mHPSE^h8^, hHPSE^h8^, m50K^hN^, m50K^hM^, or m50K^hC^ (see “Cloning” for more details on the constructs used). At 48 hr post transfection, cellular localization of HPSE was confirmed using the technique described above (see “Cellular fractionation”). To determine the antiretroviral effect of the chimeric HPSE constructs, we performed the same transfection as above but also included the FMLV molecular clone. At 48 hr post transfection, cell lysate and supernatant were harvested, processed and analyzed by immunoblotting as described above (see “Western blot analysis”).

### Particle infectivity

WT and HPSE-/- (KO) MEF cells were infected with FMLV (1 MOI) via spinoculation as mentioned above (see “Restriction by endogenous mHPSE”). At 48 hr post infection, virus was collected from the media of the infected cells followed by centrifugation at 700 x g for 10 min at 4°C, filtering through a 0.45 μm filter and treatment with DNase at a concentration of 1 μg/ml for 30 min at 37°C. Virus yields were then determined by p30 (CA) ELISA using the QuickTiter MuLV Core Antigen Elisa Kit (Cell Biolabs INC, VPK-156) as per the manufacturer’s instructions. *Mus dunni* cells, which do not express mHPSE, were seeded at 5 x 10^4^ per well of a 12-well plate and the following day infected with the aforementioned purified FMLV (20ng of CA) in the presence of 2 μg/ml of polybrene by incubating the cells with virus for 2 hr at 37°C along with periodic mixing. At 24 hr post infection, DNA was isolated, and qPCR was performed as described above (See “Restriction by endogenous mHPSE”).

### MLV Integration assay

MLV envelope pseudotyped viruses carrying a luciferase reporter genome (pFB-Luc) were produced using a Lipofectamine 3000 transfection kit, as per the manufacturer’s recommendations, with 0.75 μg of PSV MLV Eco, 1.5 μg pFB-Luc, and 2.5 μg of FMLV Gag/Pol per well. At 48 hr post transfection pseudoviruses were harvested as mentioned above (see “Particle infectivity”). 293T-mCAT-1 cells were seeded in 12-well plates and the following day transfected, using a Lipofectamine 3000, with 1 μg of either mHPSE-Myc/FLAG, hHPSE- V5/His, or EV. At 24 hr post transfection, cells were infected with the MLV envelope pseudotyped viruses mentioned above in the presence of 2 μg/ml polybrene for 2 hr at 37°C with periodic mixing along with or without Raltegravir (100nM, NIH-ARP, 11680). Cells were passaged for 6 days to eliminate episomal DNA as previously done [45]. DNA was isolated and qPCR was performed as described above (see “Restriction by endogenous mHPSE”) with primers targeting *f.luciferase* (5’-CGGAAAGACGATGACGGAAA-3’/5’- CGGTACTTCGTCCACAAACA-3’) and *hGAPDH* (5’- AACGGGAAGCTTGTCATCAATGGAAA-3’/5’-GCATCAGCAGAGGGGGCAGAG-3’).

### Effect of HPSE on MLV RNA levels

293T cells were seeded at 5 x 10^5^ per well in a 6-well plate and the following day were co- transfected using Lipofectamine 3000, as per the manufacturer’s recommendations, with 5.8 μg of FMLV and either 1 μg of hHPSE-V5/His, 1 μg of EV, or 1, 0.5, 0.1, or 0.05 μg of mHPSE- Myc/FLAG. At 48 hr post transfection, cells were lysed, RNA was isolated using an RNeasy Mini Kit followed by treatment with DNase as per the manufacturer’s instruction. cDNA was generated using SuperScript III First-Strand Synthesis System for RT-PCR per manufacturer’s recommendations. RT-qPCR was then performed on a CFX384 Touch Real-Time PCR Detection System using PowerUp SYBR Green Master Mix and primers targeting MLV U3 (5’-GCCCTCAGATGCTGCATATAA-3’/5’-TTTTTTTTTTTTTTTTTTTTTTTTTTGAAG-3’), MLV *env* (5’-TACAGGGAGCTTACCAGGCA-3’/5’-GTTCCTATGCAGAGTCCCCG-3’), MLV U5-*gag* (5’-GCCTCAATAAAGCTTGCCTTGA-3’/5’-GGGCGCCACTGCTAGAGA- 3’), and *gapdh* (5’-AACGGGAAGCTTGTCATCAATGGAAA-3’/5’- GCATCAGCAGAGGGGGCAGAG-3’).

### Effect of endogenous mHPSE on MLV

WT and HPSE-/- (KO) MEF cells were seeded at 3 x 10^4^ cells/well in 12-well plates and infected with MLV envelope pseudotyped viruses carrying a luciferase reporter genome as described above (see “Integration of MLV”). Cells were lysed 48 hr later and luciferase levels were measured as described above (see “Luminescence assays”). In parallel, additional transfected cells were passaged for 6 dpi and MLV integration was measured as described above (see “Integration of MLV”).

### RNA rate of decay

293T cells were seeded at 5 x 10^5^ cells per well in a 6-well plate and the following day co- transfected using a Lipofectamine 3000 transfection kit, as per the manufacturer’s recommendations, with 5.8 μg of FMLV and 1 μg of either mHPSE-Myc/FLAG or EV. At 48 hr post transfection, cells were treated with 20 μg/ml of Actinomycin D (Invitrogen, A7592). RNA was isolated at 0, 8, or 24 hr post drug treatment and 400 ng of purified RNA was used for cDNA synthesis followed by RT-PCR using primers defined above (see “Effect of HPSE on MLV RNA levels”).

### RNA export assays

293T cells were seeded at 5 x 10^5^ per well in a 6-well plate. Next day, cells were co-transfected using Lipofectamine 3000, as per the manufacturer’s recommendations, with 3.5 μg of FMLV and 1 μg of either mHPSE or EV. RNA and protein lysates were isolated 48 hr later from nuclear and cytoplasmic fractions using the PARIS^TM^ kit (Invitrogen, AM1921) as per the manufacturer’s instructions. Cell lysates were processed by western blots and probed for GAPDH, as described above (see “Western blot analysis”), to determine nuclear fraction purity. RNA samples were used for RT-qPCR analysis as described above using hGAPDH primers (see “Effect of HPSE on F-MLV RNA levels”), and primers targeting the MLV splice junction (nts 3347-3597) (5’-GATATCGGGCCTCGGCCAAG-3’/5’-AAACAGAGTCCCCGTTTTGG-3’) to detect spliced MLV RNA products.

### RNA splicing assays

293T cells were seeded at 5 x 10^5^ per well in a 6-well plate and co-transfected using a Lipofectamine 3000, as per the manufacturer’s recommendations, with 5.8 μg of FMLV and 1 μg of either mHPSE or EV. At 48 hr later, cells were lysed followed by RNA isolation and DNase treatment. 1 μg of RNA was used as a template for cDNA synthesis, and RT-qPCR was performed as above (see “Effect of HPSE on MLV RNA levels”) with primers targeting the MLV splice junction (see “RNA export assays**”**) and primers for the detection of unspliced MLV genome (5’-CGTGGTCTCGCTGTTCCTTGG-3’/5’-GCGGACCCACACTGTGTC-3’).

### Promoter-specific restriction

293T cells were seeded at 7 x 10^5^ per well in a 6-well plate followed by co-transfections using a Lipofectamine 3000, as per the manufacturer’s recommendations, with 3.5 μg of either CMV IE, TK, β-actin-Luc, MMLV-LTR-Luc or HIV-1 LTR-Luc and 1 μg of either EV or mHPSE- Myc/FLAG. After 48 hr, luciferase expression was measured as described above (see “Luminescence assay”). Alternatively, NIH3T3 cells were seeded at 2.5 x 10^5^ per well in a 6- well plate followed by co-transfections as described above and luciferase measurements (see “Luminescence assays”). Similarly, WT and HPSE-/- (KO) MEF cells were seeded at 5 x 10^5^ per well in a 6-well plate followed by transfections as shown above and 24 hr later luciferase expression was determined as previously described (see “Luminescence assays”).

### Evaluation of mutations in transcription factor binding sites

293T cells were seeded at 5 x 10^5^ per well in a 6-well plate followed by co-transfections using Lipofectamine 3000, as per the manufacturer’s recommendations, with 3.5 μg of either HIV-1 LTR-Luc, HIV-1 LTR-Luc constructs containing mutations in SP1, NFκB/NFAT, USF, TCF1α, or NF-IL6 binding sites, MMLV-LTR-Luc, or MMLV-LTR-Luc constructs containing mutations in SP1, or CBF/Runx binding sites and 1 μg of EV or mHPSE-Myc/FLAG. At 48 hr post transfection, luciferase expression was measured as described above (see “Luminescence assays”). Additionally, 293T cells were seeded at 5 x 10^5^ per well in a 6-well plate followed by co-transfections using Lipofectamine 3000, as per the manufacturer’s recommendations, 500ng of HIV-1 LTR-Luc constructs containing mutations in SP1, NFκB/NFAT, USF, TCF1α or NF- IL6 binding sites, and 3 μg of either EV or mHPSE-Myc/FLAG along with Tat [46]. At 48 hr post transfection, luciferase expression was measured as described above (see “Luminescence assays”). Alternatively, WT and HPSE-/- (KO) MEF cells were seeded at 5 x 10^5^ per well in a 6- well plate followed by transfections using Lipofectamine 3000, as per the manufacturer’s recommendations, with either 4.5 μg of MMLV-LTR-Luc, 4.5 μg MMLV-LTR-Luc with mutations in CBF/Runx binding sites, 10 μg MMLV-LTR-Luc with mutations in SP1 binding sites, or 1 μg of β-actin-Luc, and 24 hr post transfection, luciferase expression was measured as described above (see “Luminescence assays”).

### DNA pulldown

WT and HPSE-/- (KO) MEF cells were seeded at 1.75 x 10^6^ cells per plate in 10cm plates and the following day were infected with equal volumes of MLV envelope pseudotyped viruses carrying a luciferase reporter genome (pFB-Luc). Pseudovirus production and MEF infections were performed as described above (see “Integration of MLV” and “Effect of endogenous mHPSE on MLV”). At 48 hr post infection cells were harvested and DNA immunoprecipitation was performed as previously described with minor changes [45, 89]. Briefly, cells were initially fixed in PBS containing 1% formaldehyde (Thermo Scientific), crosslinking was halted by the addition of glycine and cells were then collected in nuclear lysis buffer (1% SDS, 0.01M EDTA, and 0.05M pH 8 Tris-HCl). Samples containing 10 μg of DNA were then incubated overnight at 4°C with either anti-RNA pol II antibody (mouse anti-Rpb1 CTD, Cell Signaling, 2629S), mouse anti-SP1 antibody (Santa Cruz Biotechnology, sc-17824X), or mouse IgG (Invitrogen, 31878) in ChIP dilution buffer (0.01% SDS, 1.1% Triton X-100, 1.2mM EDTA, 16.7mM pH 8 Tris-HCl, and 0.167M NaCl). The following day, a mixture of 12.5 μl protein A (Invitrogen, 10001) and 12.5 μl protein G (Invitrogen, 10003) Dynabeads pre-incubated in ChIP dilution buffer supplemented with glycogen (Invitrogen, AM9510), yeast RNA (Invitrogen, AM7118), and recombinant albumin (New England BioLabs,. B9200S) was added to lysates. Samples were then incubated with Dynabeads for 2hr at 4°C and washed as previously described [89] prior to being eluted in elution buffer (1% SDS, 1mM EDTA, 0.01M pH 8 Tris-HCl, and 0.2M NaCl) supplemented with proteinase K (Qiagen, 1114885) and Rnase A (Invitrogen, no.8003089) for 30 min at 65°C. Crosslinking was then reversed in eluted samples in a 65°C incubation overnight. DNA was isolated using a QIAquick PCR Purification Kit (Qiagen, 28106) according to manufacturer’s instructions. Equal volumes of immunoprecipitated samples along with input samples, collected prior to immunoprecipitation, were used to perform qPCR with primers targeting the MLV LTR (5’-CGCTGACGGGTAGTCAATCACT-3’/5’- GTCCAGCCCTCAGCAGTTTC-3’).

### Co-immunoprecipitation**s**

293T cells were seeded at 5 x 10^5^ per well in a 6-well plate and the following day co-transfected using a Lipofectamine 3000, as per the manufacturer’s recommendations, with 5 μg of mSP1 and 100ng of either mHPSE-V5/His, hHPSE-V5/His, or pCDNA. Both mHPSE and hHPSE-V5/His had N-terminal Myc tag additions and are described above (see “Cloning”). At 48 hr post transfection, cells were collected in PBS followed by centrifugation at 300 x g for 10 min at 4°C. Cell pellets were then resuspended in NP-40 Lysis Buffer (Research Products International, N32000) supplemented with 2X Halt Protease and Phosphatase Inhibitor Cocktail (Thermo Scientific, 78442), and indicated samples were treated with 750 U/ml benzonase (Sigma-Aldrich, E1014). 50 μl of protein A Dynabeads were first incubated with either mouse anti-Myc antibody (Cell Signaling Technology, 2276S) or mouse IgG for 15 min at room temperature. Beads were then incubated with 2.5 mg of cell lysate for 2 hr at room temperature, washed and protein complexes were collected in elution buffer (Invitrogen). Additionally, 10μl of each sample was resolved on a 1% agarose gel.

### SP1-mediated enhancement of MLV production

293T cells were seeded at 5 x 10^5^ per well in a 6-well plate followed by co-transfections using Lipofectamine 3000, as per the manufacturer’s recommendations, with increasing amounts (1 μg, 3 μg, or 5 μg) of mSP1, 3.5 μg of FMLV, 1 μg of pFB-Luc, and 50 ng of either mHPSE- Myc/FLAG or EV. Virus produced was harvested after 48 hr followed by centrifugation at 700 g for 10 min at 4°C and passing it through at 0.45 μm filter. To determine virus yield, equal volumes of virus were then used to infect NIH3T3 cells by spinoculation followed by measurement of luciferase levels as described above (see “Restriction by endogenous mHPSE,” and “Luminescence assays”).

### Effect of mHPSE on SP1 degradation

NIH3T3 cells were seeded in a 6-well plate at 3.5 x 10^5^ cells per well and transfected 24 hr later in serum free DMEM using Lipofectamine LTX, as per the manufacturer’s recommendations, with 0.02, 0.1, 0.5, 1 or 2.5 μg of mHPSE-Myc/FLAG or EV. Cell lysates were harvested 48 hr after transfection and western blots were performed to determine endogenous mSP1 and mHPSE-Myc/FLAG levels as described above (see “Western blot analysis”). In the case of 293T cells, cells were seeded at 4.5 x 10^5^ in a 6-well plate followed by transfections using Lipofectamine 3000, as per the manufacturer’s recommendations plate, with 0.02, 0.1, 0.5, 1, or 2.5 μg of mHPSE or EV. Cell lysates were harvested 48 hr later and western blots were performed to determine endogenous hSP1 and mHPSE levels as described above (see “Western blot analysis”).

### HPSE Enzymatic Activity Assay and Restriction

The enzymatic activity of HPSE was measured using a Heparan Degrading Enzyme Assay Kit (Takara, MK412) as per the manufacturer’s recommendations. In brief, samples were initially incubated with biotinylated heparan sulfate. The mixture was then transferred to a plate coated with fibroblast growth factor (FGF). Plates were then washed followed by the addition of POD-conjugated avidin. After 5 min, absorbance was measured using a Biostack4 luminometer. To evaluate the effect of SF4 (DiagnoCine, FNK-11829) on the antiretroviral activity of mHPSE, 293T cells were seeded at 0.25 x 10^5^ per well in a 96-well plate followed by co-transfections using Lipofectamine 3000, as per the manufacturer’s recommendations, with 50ng of MLV-LTR-Luc and 10ng of either EV or mHPSE-Myc/FLAG. At 6 hr post transfection, cells were treated with 100 μM of either DMSO or SF4. After an additional 24 hr, luciferase expression was quantified as described above (see “Luminescence assays”) and, in parallel, enzymatic activity was measured using the kit mentioned in this section. Additionally, WT MEFs were seeded at 5 x 10^5^ per well in a 6-well plate followed by transfection using Lipofectamine 3000, as per the manufacturer’s recommendations, with 4.5 μg of either MLV-LTR-Luc or HIV-1 LTR-Luc. SF4 or DMSO treatment and measurement of luciferase expression were performed as described with 293T cells. To evaluate the effect of OGT 2115 (Tocris, 2710) on the antiretroviral effect of mHPSE, 293T cells were seeded at 5 x 10^5^ per well in a 6-well plate followed by co-transfections using Lipofectamine 3000, as per the manufacturer’s recommendations, with 3.5 μg of MLV-LTR-Luc and 1 μg of either EV or mHPSE-Myc/FLAG. At 12 hr post transfection, cells were treated with 10 μM of either DMSO or OGT 2115 [19] and 48 hr post transfection luciferase expression was quantified as described above (see “Luminescence assays”). To determine the effect of an enzymatically inactive mHPSE or a mHPSE with defects in HS binding, 293T cells were seeded at 7 x 10^5^ per well in a 6-well plate followed by co-transfections using Lipofectamine 3000, as per the manufacturer’s recommendations, with 3.5 μg of either MMLV-LTR-Luc or HIV-1 LTR-Luc and 1 μg of either EV, mHPSE-Myc/FLAG, mHPSE EE217/335AA or mHPSE KK150/153AA. At 48 hr post transfection, enzymatic activity was measured as above. Luciferase expression was quantified as previously described (see “Luminescence assays”) and cellular localization was determined as shown in a previous section (see “Cellular fractionation”).

### Effect of Akt pathway inhibition on HPSE-mediated MLV restriction

293T cells were seeded at 5 x 10^5^ per well in a 6-well plate and the following day treated with 10 μM DMSO or LY 294002 (Sigma-Aldrich, 440204). At 4hr post treatment, cells were co- transfected using Lipofectamine 3000, as per the manufacturer’s recommendations, with 3.5 μg of MLV-LTR-Luc and 1 μg of either EV or mHPSE. At 20 hr post transfection, Akt and p-Akt levels were analyzed by western blot analysis as described above (see “Western blot analysis”) and luciferase expression was quantified as previously discussed (see “Luminescence assay”). IGF1-FLAG (Addgene, 175170), a known activator of the AKT pathway [60], served as a positive control.

### Statistical analyses

Statistical analyses were performed using GraphPad Prism software version 10.0. The statistical tests used are described in the manuscript. A difference was considered to be significant for P values of 0.05.

## Acknowledgments

We thank members of the Stavrou lab for insightful discussions on this project. We thank Stephen Goff, Gary Wang for providing us with protocols for the DNA pulldown experiments of the project. We also thank Susan Ross, Christine Kozak, Leonard “Pug” Evans (deceased), Alan Rein and Frank Kirchhoff for reagents. This work was supported by SUNY University at Buffalo startup funds.

---

**Supplementary Figure S1.**
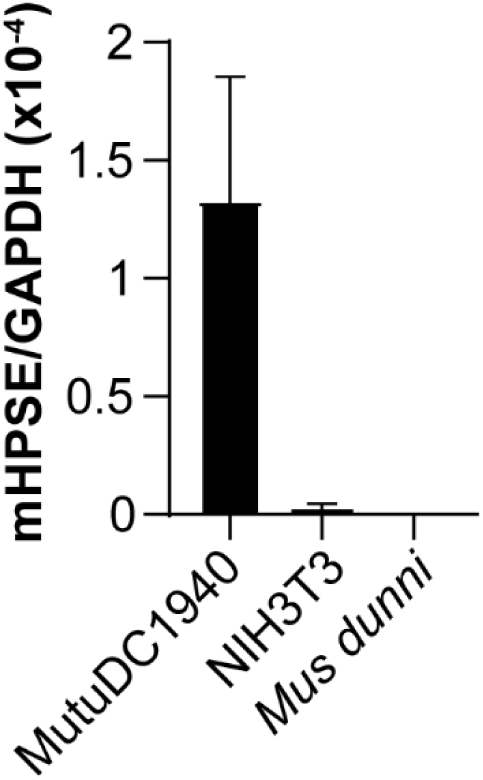
Mouse *Hpse* is not expressed in NIH3T3 or *Mus dunni* cells. RNA was isolated from MutuDC1940, NIH3T3, and *Mus dunni* cells, followed by RT-qPCR to evaluate *Hpse* expression levels normalized to *Gapdh*. (N=3)

**Supplementary Figure S2.**
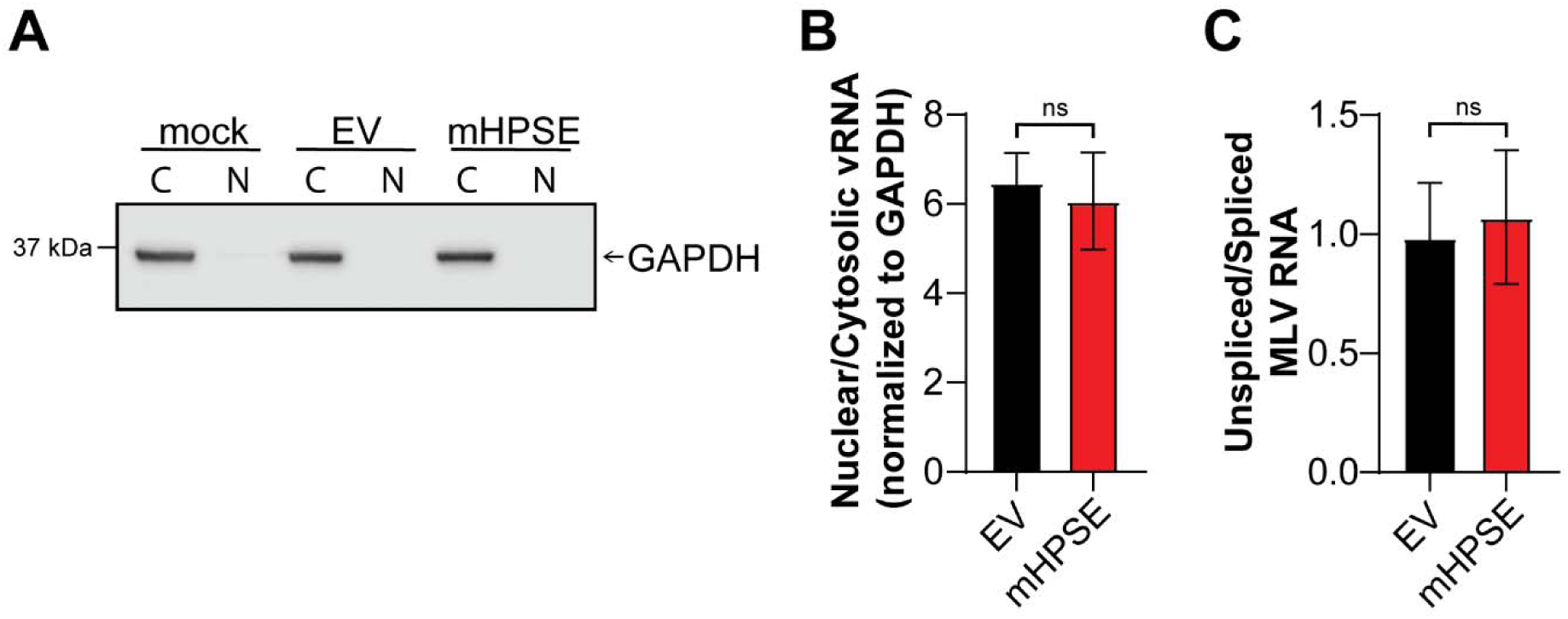
Mouse heparanase (mHPSE) does not affect the export or splicing of MLV RNA. 293T cells were co-transfected with plasmids encoding an MLV infectious clone and either mHPSE or empty vector (EV). **(A-B)** mHPSE does not affect MLV RNA export. Following co-transfection, samples were fractionated into cytosolic (C) and nuclear (N) fractions. **(A)** Purity of nuclear fractions was analyzed by immunoblotting for the cytosolic protein, GAPDH. Representative western blot of 3 independent experiments is shown. **(B)** MLV RNA transcripts in C and N fractions were quantified by RT-qPCR. Data presented as the ratio of MLV RNA in N/C fractions in the presence of mHPSE or EV and normalized to *GADPH*. **(C)** mHPSE does not affect the splicing of MLV RNA. Following co-transfection, levels of spliced and unspliced MLV RNA were determined by RT-PCR and the ratio of unspliced/spliced MLV RNA was calculated. All data in graphs presented as mean ± SEM. Statistical analysis was performed by unpaired t test, N=3, ns; not significant.

**Supplementary Figure S3.**
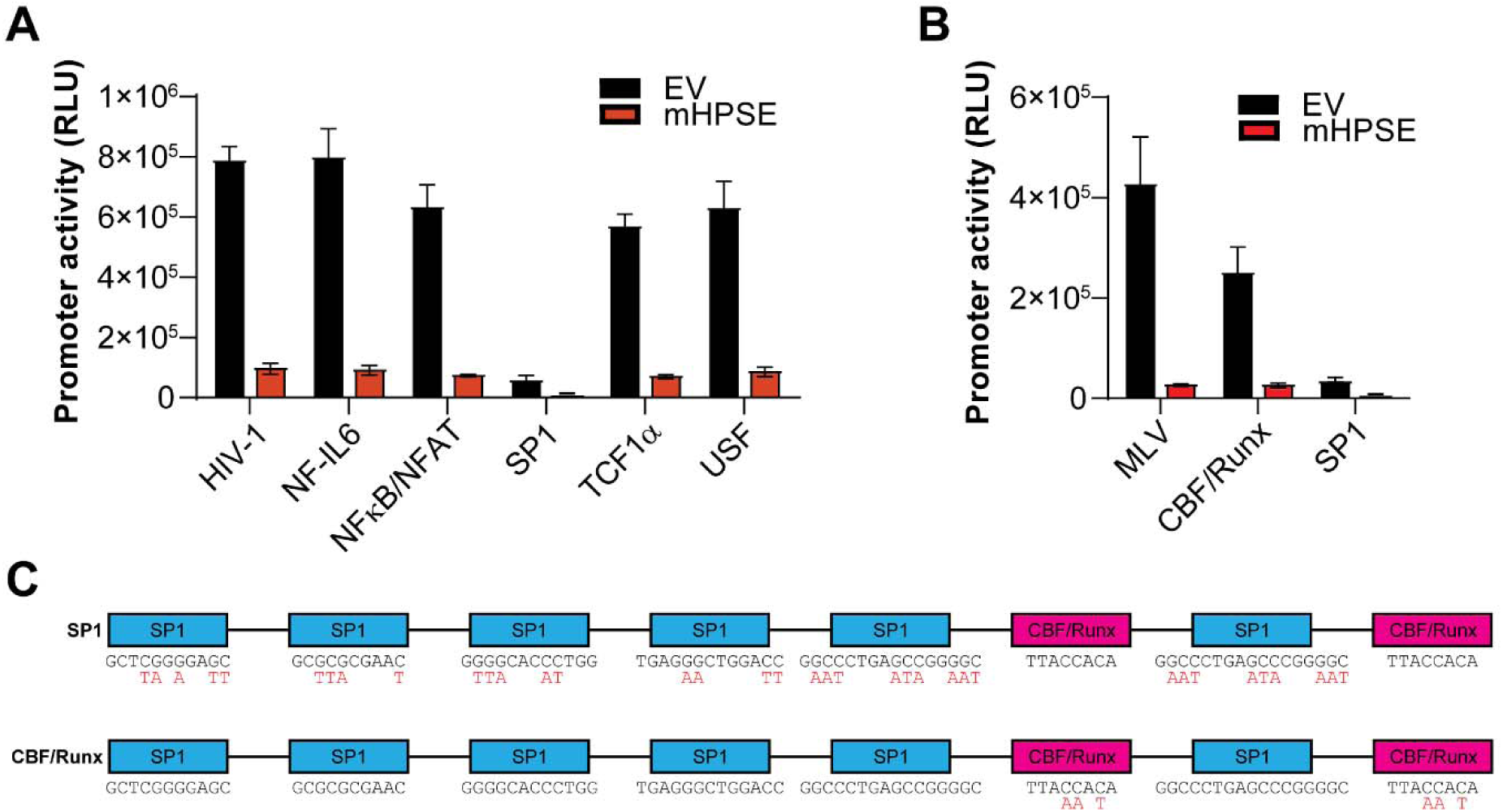
Mutations in SP1 binding sites reduce retrovirus LTR activity. 293T cells were co-transfected with **(A)** HIV-1 or **(B)** MLV LTR luciferase reporter constructs containing mutations at the indicated transcription factor binding site along with plasmids expressing either EV or mouse heparanase (mHPSE). Data presented as mean ± SEM, **(A)** N=4 and **(B)** N=3, RLU; relative light units. **(C)** Diagram displaying the mutations made in the SP1 or CBF/Runx binding sites of the MLV LTR.

**Supplementary Figure S4.**
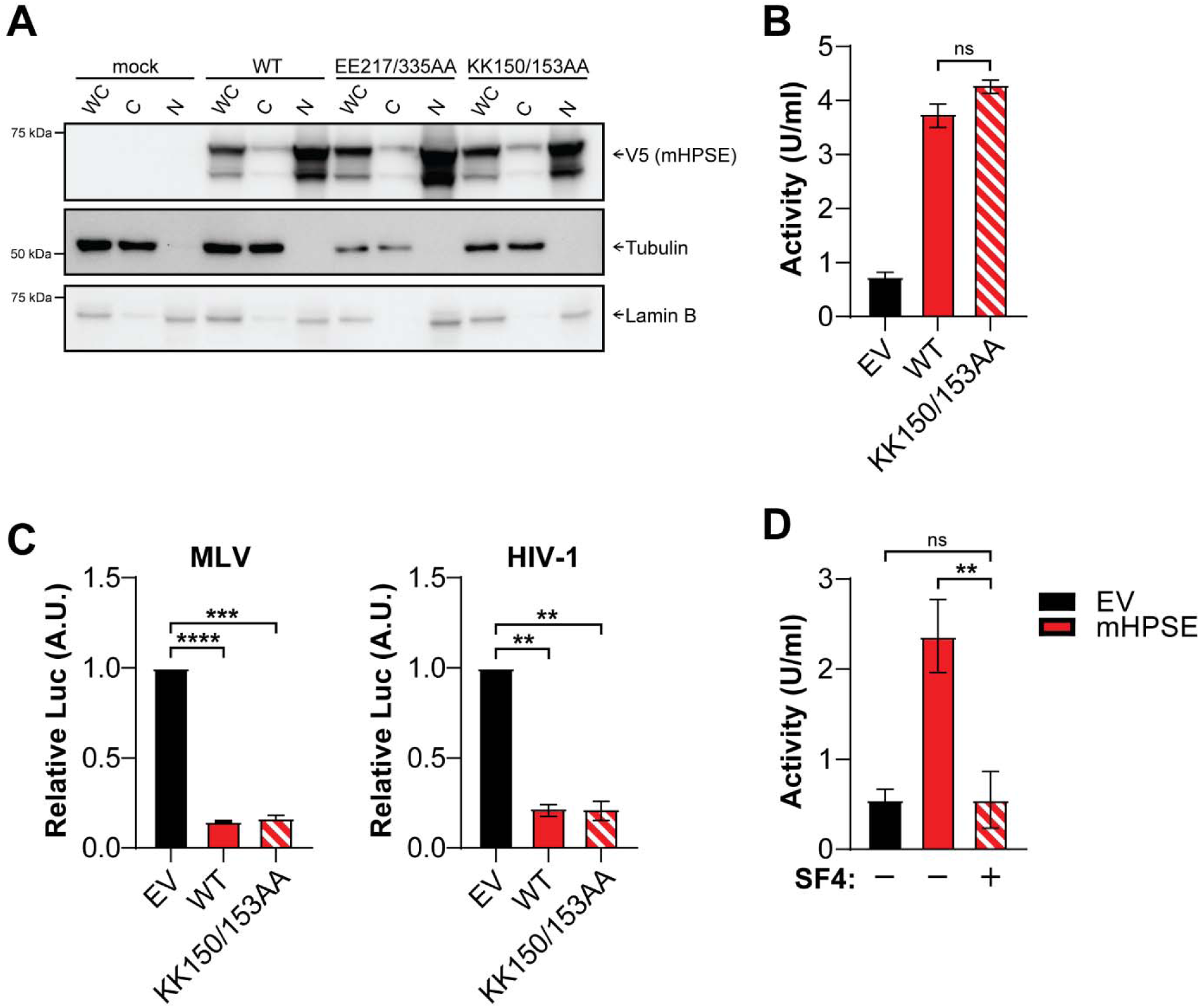
KK150/153 of mouse heparanase (mHPSE) do not affect its antiretroviral function. **(A)** mHPSE (WT) glutamic acid (E) residues 217 and 335 and lysine (K) residues 150 and 153 do not affect mHPSE nuclear localization. Cells transfected with the indicated plasmids were fractionated into whole cell lysate (WC), cytosolic fraction (C), or nuclear fraction (N) followed by immunoblotting. Immunoblots shown are representative of 3 independent experiments. (**B**) K residues 150 and 153 do not affect the enzymatic activity of mHPSE. 293T cells were transfected with plasmids encoding either empty vector (EV), wild type mHPSE (WT), or mHPSE in which either K residues 150 and 153 are substituted for alanine (KK150/153AA) followed by measurement of mHPSE enzymatic activity. **(C)** Lysine residues 150 and 153 are dispensable to the antiretroviral function of mHPSE. 293T cells were co- transfected with either MLV LTR or HIV-1 LTR luciferase reporter constructs and plasmids encoding EV, WT, or KK150/153AA followed by luciferase measurements. **(D)** SF4 inhibits the enzymatic activity of mHPSE. 293T cells were transfected with either EV or mHPSE and then treated with SF4 (100 μM) or vehicle control, followed by measurement of enzymatic activity. All data in graphs presented as mean ± SEM. Statistical analysis was performed by **(B)** unpaired t test, N=4, **(C)** one sample t test, N=3, or **(D)** one-way ANOVA, Dunnett’s multiple comparison test, ns; not significant, \*\**P*≤0.01, \*\*\**P*≤0.001, \*\*\*\**P*≤0.0001.

**Supplementary Figure S5.**
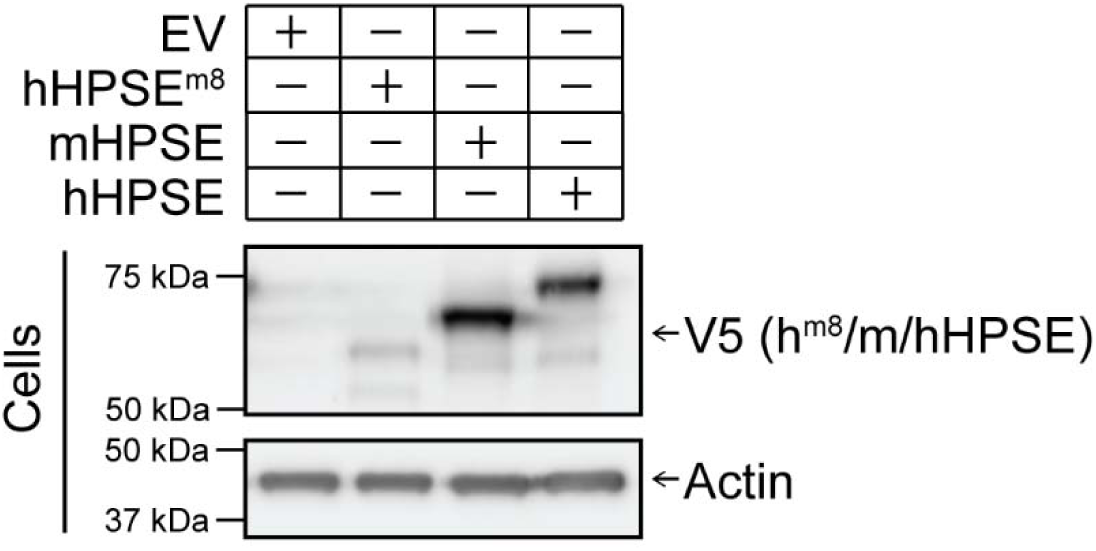
hHPSE^m8^ expresses at very low levels. 293T cells were transfected with plasmids encoding either empty vector (EV), mouse heparanase (mHPSE), human heparanase (hHPSE), or hHPSE^m8^, and protein samples were analyzed by immunoblotting for the indicated proteins. Displayed immunoblots are representative of 3 independent experiments.

**Supplementary Table S1:**
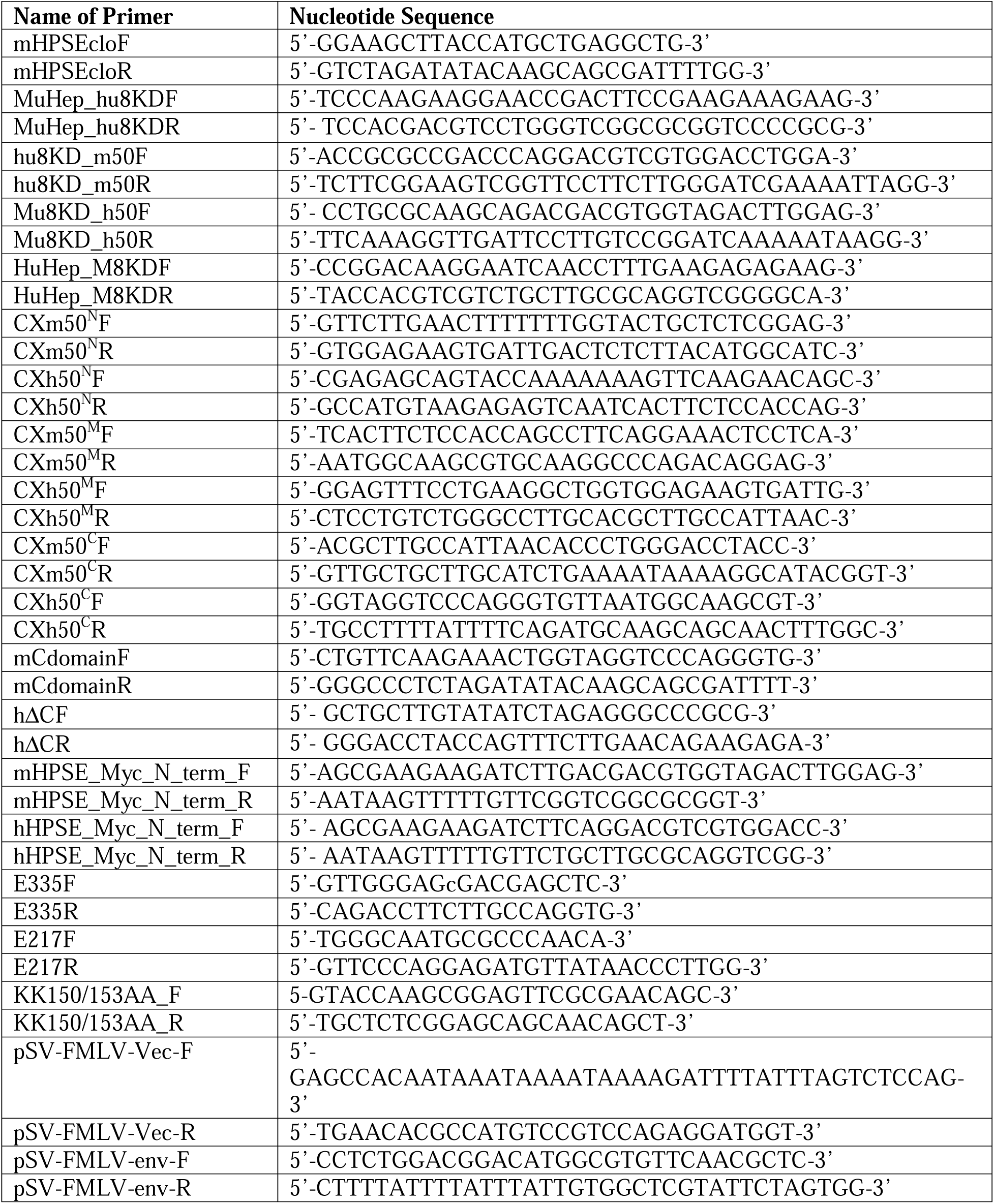
Primers.

